# Comprehensive profiling of small RNAs and their changes and linkages to mRNAs in schizophrenia and bipolar disorder

**DOI:** 10.1101/2024.12.24.630254

**Authors:** Stepan Nersisyan, Phillipe Loher, Iliza Nazeraj, Zhiping Shao, John F. Fullard, Georgios Voloudakis, Kiran Girdhar, Panos Roussos, Isidore Rigoutsos

**Affiliations:** Computational Medicine Center, Thomas Jefferson University, Philadelphia, PA, USA; Center for Disease Neurogenomics, Icahn School of Medicine at Mount Sinai, New York, USA; Friedman Brain Institute, Icahn School of Medicine at Mount Sinai, New York, USA; Department of Psychiatry, Icahn School of Medicine at Mount Sinai, New York, USA; Department of Genetics and Genomic Sciences, Icahn School of Medicine at Mount Sinai, New York, USA; Center for Precision Medicine and Translational Therapeutics, JJ Peters VA Medical Center, Bronx, New York, USA; Mental Illness Research Education and Clinical Center (MIRECC), JJ Peters VA Medical Center, Bronx, New York, USA

**Keywords:** schizophrenia, bipolar disorder, brains, small non-coding RNAs, miRNA, miRNA isoforms, isomiRs, tRNA-derived fragments, tRFs, rRNA-derived fragments, rRFs, Y RNA-derived fragments, yRFs

## Abstract

We investigated small non-coding RNAs (sncRNAs) from the prefrontal cortex of 93 individuals diagnosed with schizophrenia (SCZ) or bipolar disorder (BD) and 77 controls. We uncovered recurring complex sncRNA profiles, with 98% of all sncRNAs being accounted for by miRNA isoforms (60.6%), tRNA-derived fragments (17.8%), rRNA-derived fragments (11.4%), and Y RNA-derived fragments (8.3%). In SCZ, 15% of all sncRNAs exhibit statistically significant changes in their abundance. In BD, the fold changes (FCs) are highly correlated with those in SCZ but less acute. Non-templated nucleotide additions to the 3′-ends of many miRNA isoforms determine their FC independently of miRNA identity or genomic locus of origin. In both SCZ and BD, disease- and age-associated sncRNAs and mRNAs reveal accelerated aging. Co-expression modules between sncRNAs and mRNAs align with the polarities of SCZ changes and implicate sncRNAs in critical processes, including synaptic signaling, neurogenesis, memory, behavior, and cognition.

## INTRODUCTION

Schizophrenia (SCZ) and bipolar disorder (BD) are complex neuropsychiatric disorders that significantly impact cognitive and emotional functioning. Transcriptomic studies of post-mortem brain tissues have been instrumental, highlighting alterations in gene expression in affected brains, particularly in pathways associated with synaptic signaling, neurogenesis, and neurotransmitter systems, thus providing insights into the molecular underpinnings of SCZ and BD [1, 2]. Additionally, genome-wide association studies (GWAS) have identified numerous risk loci associated with these disorders, underscoring the polygenic nature of their etiology [3]. Despite these advances, the heritability explained by GWAS is limited, suggesting the potential involvement of non-coding genes and other genomic regions.

Small non-coding RNAs (sncRNAs) are regulatory in nature and represent a sizable component of an organism’s transcriptome [4]. Even though they outnumber protein-coding genes by two orders of magnitude, sncRNAs have remained largely uncharacterized outside the cancer context [5]. Among the various classes, the best-studied sncRNA species are microRNAs (miRNAs), 18-22 nucleotides (nts) in length, that bind their mRNA or long non-coding RNA targets in a sequence-dependent manner [4, 6].

Several previous efforts in the SCZ context relied on miRNA microarray or PCR array analyses to identify differentially abundant (DA) miRNAs between post-mortem brain samples of cases and controls [7–13]. For more information, the reader is referred to several excellent recent reviews on miRNAs in SCZ and other psychiatric disorders [14, 15]. The results of these studies showed limiter concordance, presumably resulting from combinations of factors known to affect differential miRNA expression analysis, including differences in miRNA profiling platforms [16], batch effects [17], differing clinical and demographic variables (e.g., ethnicity, sex, age) [18, 19], differences in the analyzed brain regions [20]. Nowadays, RNA-seq has superseded these earlier approaches, offering improved sensitivity, specificity, and the possibility of data-driven normalization [21–23]. Additionally, RNA-seq provides a unique opportunity for the unbiased characterization of sncRNAs in a sample.

One of the early successes of RNA-seq was the discovery in 2008 of miRNA isoforms (isomiRs) [24]. IsomiRs are co-expressed mature miRNAs that arise from the same miRNA arm and have distinct sequences [4]. IsomiRs typically differ from one another and the “reference” sequences found in public databases, such as miRBase [25], by a few nts at either their 5′-end or 3′-end. IsomiRs whose sequences match the human genome exactly are categorized as “templated.” The primary source of templated isomiRs is the systematic and regulated heterogeneous cleavage of miRNA precursor hairpins by DROSHA and DICER enzymes [26]. The “non-templated” isomiRs are another emerging rich category: these are produced through the post-transcriptional addition of nucleotides to their 3′-ends [26, 27]. Sibling isomiRs that differ even by a single nucleotide, whether templated or non-templated, can target different genes [26, 28, 29] and exhibit different sub-cellular localization [30].

The universe of regulatory sncRNAs includes additional molecular classes, namely, the tRNA-derived fragments (tRFs), the rRNA-derived fragments (rRFs), and the Y RNA-derived fragments (yRFs) [4, 31]. We previously showed that the abundance levels of isomiRs, tRFs, rRFs, and yRFs depend on “context” (e.g., tissue type, disease) [31–33], as well as “personal attributes” (e.g., ancestry and sex) [18, 34, 35]. Despite their omnipresence, high abundance, and intriguing dependencies, tRFs, rRFs, and yRFs are less studied compared to miRNAs/isomiRs.

In this study, we comprehensively characterized sncRNAs in post-mortem brain samples of SCZ and BD cases and controls. We also identified those sncRNAs that are DA between cases and controls and linked them through co-expression analysis to protein-coding genes from several critical pathways. We ensured robustness by including 170 brain samples from two independent brain banks.

## MATERIALS AND METHODS

### Description of post-mortem brain samples

Frozen brain tissue from the prefrontal cortex (PFC, Brodmann areas 9 and 46) was obtained from two brain banks, as detailed below. This cohort includes 170 SCZ, BD, and control subjects (n = 53, 40, and 77, respectively), selected based on strict inclusion/exclusion criteria. All subjects met the relevant DSM-IV diagnostic criteria, determined through consensus conferences involving a review of medical records, direct clinical assessments, and interviews with family members or caregivers. Tissue donors were sourced from the Icahn School of Medicine at Mount Sinai (MSSM) and the NIMH Human Brain Collection Core (HBCC). Supp. Table S1 provides sample-level demographic information, including sex, age at death, post-mortem interval (PMI), and stratification by institution and diagnosis. All included subjects are of Caucasian ancestry.

### RNA isolation, library preparation, and sequencing

We isolated RNA using miRNeasy kits (Qiagen) according to the manufacturer’s instructions. RNA integrity number (RIN) was assessed using a 2200 TapeStation (Agilent Technologies). All samples had RIN ≥ 6 (Supp. Table S1). We prepared sequencing libraries using NEBNext Small RNA Prep Set for Illumina (NEB #E7330) at the Jefferson Genomics Core Facility according to the standard kit protocols, which size-selected for sncRNAs. The NEBNext 3′-adapter used is AGATCGGAAGAGCACACGTCT. We sequenced all samples on the same Illumina NextSeq 500 sequencing platform at 75 cycles. We used cutadapt v4.6 [36] to remove adapters and low-quality bases from the 3′ ends of sequenced reads, discarding reads that contained no adapter or were shorter than 16 nucleotides with (cutadapt --match-read-wildcards -q 15 -e 0.12 -a AGATCGGAAGAGCACACGTCT -m 16 --discard-untrimmed).

### SncRNA-seq read mapping

We used IsoMiRmap [37] and MINTmap [38] to profile isomiRs and tRFs. We used the previously described exhaustive, brute-force search [31] to profile rRFs, yRFs, and other repetitive fragments (rpFs) by mapping the reads to the reference rRNAs [35, 39], Y RNAs [31], and other repetitive elements annotated by RepeatMasker [40], respectively. In all these steps, we required exact matching of reads to the universe of known isomiRs, tRFs, rRFs, yRFs, and rpFs, while providing for the post-transcriptional addition of *non-templated* nucleotides to isomiRs (e.g., 3′ uridylation or adenylation) and tRFs (3′ addition of CCA to mature tRNAs).

Next, for each sample, we identified unique reads that could not be mapped with the above process and identified those with Levenshtein distance (LD) ≤ 2 from at least one of the sample’s 10,000 most abundant annotated sncRNAs (i.e., isomiRs, tRFs, rRFs, yRFs, and rpFs). The matching reads received the prefix “unk-” (unknown), and the links between these sequences and corresponding annotated sncRNAs with LD ≤ 2 were stored in a separate field. We sought LD matches using the polyleven Python package (https://github.com/fujimotos/polyleven). Finally, we used bowtie v1.3.1 [41] to place any remaining unmapped reads to the reference GRCh38.p14 genome, allowing up to 1 replacement but no insertions or deletions (bowtie -v 1 -a --best –strata -S). We prefixed reads from this last group that matched with “bwt-” and annotated them by intersecting their coordinates with GENCODE v42’s basic annotation file using bedtools v2.30 [42].

We converted raw read counts to reads-per-million (RPM) units for filtering purposes. The analyses described below used only sncRNAs whose abundance was at least 10 RPM in at least 25% of all (case and control) samples.

### Labeling of sncRNAs

To uniformly label sncRNAs across these diverse molecule types, we used the “license plates” scheme that we originally introduced for tRFs [43, 44] and have since expanded to rRFs [35, 39], isomiRs [37], yRFs, and repetitive elements [31]. In a nutshell, the license plate of a sncRNA is a base-32 encoding of the sncRNA’s sequence prefixed by a three-letter abbreviation of the class to which the sncRNA belongs (e.g., “iso-,” “tRF-,” rRF-,” yRF-,” or “rpF-”). License plates have several desirable properties: (1) they are invertible (each sncRNA has a unique license plate and *vice versa*); (2) they obviate the need for a centralized broker group or organization that issues labels – indeed, the codes for creating license plates and converting license plates to nucleotide sequences are publicly available; and (3) they are independent of genome/transcriptome assemblies, meaning the labels persist over time.

### mRNA-seq data

We obtained transcript-level read count data for the same samples from the CommonMind Consortium [2]. We discarded samples whose RIN was less than 6, leaving us with 161 samples for mRNA analysis. We summarized transcript-level counts at the gene level using GENCODE v30 annotation. We converted raw read counts to transcripts-per-million (TPM) units for filtering purposes. The analyses described below used only mRNAs whose abundance was at least 1 TPM in at least 25% of all (case and control) samples.

### Differential abundance analysis

We used DESeq2 v1.44 to identify sncRNAs and mRNAs that are differentially abundant (DA) between cases and controls [45]. We conducted separate analyses for the HBCC and MSSM brain banks because of the acute differences in the ages of the respective donors: age at death ranges from 13 to 83 years old in the HBCC (i.e., both younger and older individuals) and from 47 to 96 in the MSSM (i.e., only older individuals). We separately normalized the counts from the HBCC and MSSM samples using standard DESeq2’s median of ratios algorithm. Then, we winsorized the normalized counts to the 5^th^ and 95^th^ percentiles of each molecule to avoid cases of outlier-driven DA results.

Before covariate selection, we assessed correlations between clinical and demographic factors (Supp. Figure S1A-B). There are statistically significant, brain bank-specific correlations primarily associated with disease status, age at death, PMI, and RIN. This indicated the need to adjust for confounders in the DA analysis. In both brain banks, RIN is negatively correlated with PMI, as expected. However, multivariable regression analysis showed a negative association of RIN with SCZ/BD that is independent of PMI (Supp. Figure S1C-D). This mirrors a similar dependence that was previously reported for post-mortem brain samples from Alzheimer’s cases [46].

We first fit univariate models testing for associations of sncRNAs and mRNAs with different known variables: disease status, age, sex, and RIN. We selected an age cutoff of 45 years to maximize the balance between cases and controls and formed two age groups based on that cutoff. We also binarized samples based on their RIN using a standard threshold of 7.5 [47, 48]. These two binarizations helped avoid assuming a linear relationship between gene expression and age or RIN. Based on the results of univariate analysis, we included disease status, age, and RIN in the multivariable model for sncRNA-seq. In addition to these variables, we added sex in the mRNA-seq model as we found DA of ∼50 genes between males and females (predominantly encoded in X and Y chromosomes).

For the HBCC brain bank, we also fit a model with age-group-specific effects of SCZ by introducing a variable generated by concatenating the labels of the disease and age groups. To ensure comparability of disease vs. control comparisons in younger and older age groups, we randomly down-sampled the younger age group to match the sample size of the older group (we used 100 down-samplings with different random seed values). Similarly, to ensure comparability of older vs. younger age group comparisons between cases and controls, we conducted separate down-samplings of the control group to match the sample size of the disease group.

We controlled for potential hidden confounding factors by estimating surrogate variables using the SVA v3.52 package [49] and adding them to the multivariable model. We generated DA results from standard contrasts (e.g., SCZ vs. control, BD vs. control) using the default Wald’s test. We also generated the DA results on the combined effect of disease (SCZ or BD) and the combined effect of disease and age (concatenated disease and age group labels) using the likelihood-ratio test. We controlled for a false discovery rate (FDR) using the Benjamini-Hochberg procedure.

### Enrichment analysis

Unlike overrepresentation analysis, gene set enrichment analysis (GSEA) does not require setting fold change (FC) or statistical significance thresholds on the DA analysis results [50]. We ran GSEA on the DESeq2-ranked lists of sncRNAs and mRNAs using fgsea v1.30 package (https://bioconductor.org/packages/release/bioc/html/fgsea.html). We downloaded reference gene sets associated with Gene Ontology (GO) terms from MSigDB v2023.2 [51]. We also used the overrepresentation analysis available in the GSEApy v1.1.3 package [52] to characterize GO terms enriched in gene co-expression clusters and miRNA target gene sets.

### Co-expression analysis in the face of confounding variables

Computing correlations between the abundances of two molecules (e.g., mRNA-mRNA or sncRNA-mRNA) can provide valuable insights into co-expression patterns and help form modules of co-expressed transcripts. While not commonly discussed, confounding variables can dramatically influence the correlation between pairs of transcripts [53]: that correlation may disappear or even change signs after confounding variables are accounted for. Supp. Figure S2 shows one such example using two genes, KDM5D and RPS4Y1, located on the Y chromosome. When we consider all samples of the HBCC brain bank, independently of their sex, the correlation between these two genes equals 0.99. On the other hand, computing the correlation using only the male samples results in a dramatically different value of -0.65: here, sex acts as a third variable and leads to an incorrect computation and a false positive call.

To reduce the effect of confounding, we implemented a two-step procedure. In the first step, we started with log_2_-transformed normalized and winsorized counts and used multivariable linear regression to residualize the effects of known factors: disease status, age, sex, and RIN. In the second step, we applied a modified instance of Parsana et al. [53] to remove hidden confounders. We iteratively regressed out principal components (PCs) from the sncRNA and mRNA expression data in a way that maximized the enrichment of known pathways among the highly correlated molecule pairs. The selected residualized PCs were PC_3_, PC_4_, PC_9_, PC_11_, and PC_13_.

The enrichment score was computed as follows. First, we labeled the top 5% of mRNA-mRNA pairs with the highest absolute Pearson correlations and the top 5% of isomiR-mRNA pairs with the most negative correlations as “highly correlated” pairs. Then, for each isomiR and mRNA, we computed a p-value (hypergeometric test) to determine whether “reference” pairs are overrepresented among the highly correlated pairs involving that specific isomiR or mRNA. We adjusted p-values using the Benjamini-Hochberg procedure and computed the global enrichment scores by averaging negative logarithms of adjusted p-values separately for isomiRs and mRNAs.

In the mRNA-mRNA case, we labeled a pair as “reference” if the corresponding two genes share at least one “Canonical Pathway” or “GO term” from MSigDB v2023.2. In the isomiR-mRNA case, “reference” indicated that the mRNA is an experimentally validated or computationally predicted target of the corresponding canonical miRNA. We restricted the analysis to isomiRs with a canonical 5′-end, as the 5′-end sequence is the primary determinant of isomiR targets [4]. We obtained a list of experimentally validated miRNA-mRNA pairs from TarBase v9.0 [54] (the main portion of the data comes from CLIP experiments). We used RNA22 v2.0 [55] to predict miRNA binding sites across the whole span of target mRNAs.

### Statistical analysis and data visualization

We used the SciPy v1.12 [56] Python module for routine statistical tests and procedures (Fisher’s exact test, Mann-Whitney’s U-test, correlation analysis, and linear regression analysis). We used the scikit-learn v1.4 (https://scikit-learn.org) Python module for PCA. We used the Seaborn v0.13 (https://seaborn.pydata.org) and Matplotlib v3.8 (https://matplotlib.org) Python modules for data visualization.

## RESULTS

### IsomiRs, tRFs, rRFs, yRFs, and other sncRNAs abound in the brains of cases and controls

We generated sncRNA sequencing data from post-mortem brain samples across two independent brain banks, which included 53 SCZ cases, 40 BD cases, and 77 controls. Using a state-of-the-art analytical pipeline (see Methods), we mapped and comprehensively annotated sequencing reads in 170 small RNA-seq datasets. First, we profiled isomiRs, tRFs, rRFs, yRFs, and rpFs, enforcing exact matching. Next, we identified those of the remaining reads with LD ≤ 2 from at least one of the sncRNAs that was annotated during the first step. Finally, we mapped all remaining reads to the human genome, allowing at most one replacement but no insertions or deletions.

In this analysis, we considered only the 7,473 mapped sncRNAs whose abundance is at least 10 RPM in at least 25% of the 170 samples. Figure 1A shows the distribution of the identified sncRNAs in each of the considered categories (an additional entry comprises unmapped molecules). Individual molecules within each sncRNA type are sorted by mean abundance, and their ranks are shown on the X-axis. The Y-axis shows the cumulative abundance (mean ± SD) of the ranked molecules. We analyzed all the samples together since the total abundance of each RNA class showed no statistically significant differences between cases and controls.

**Figure 1.**
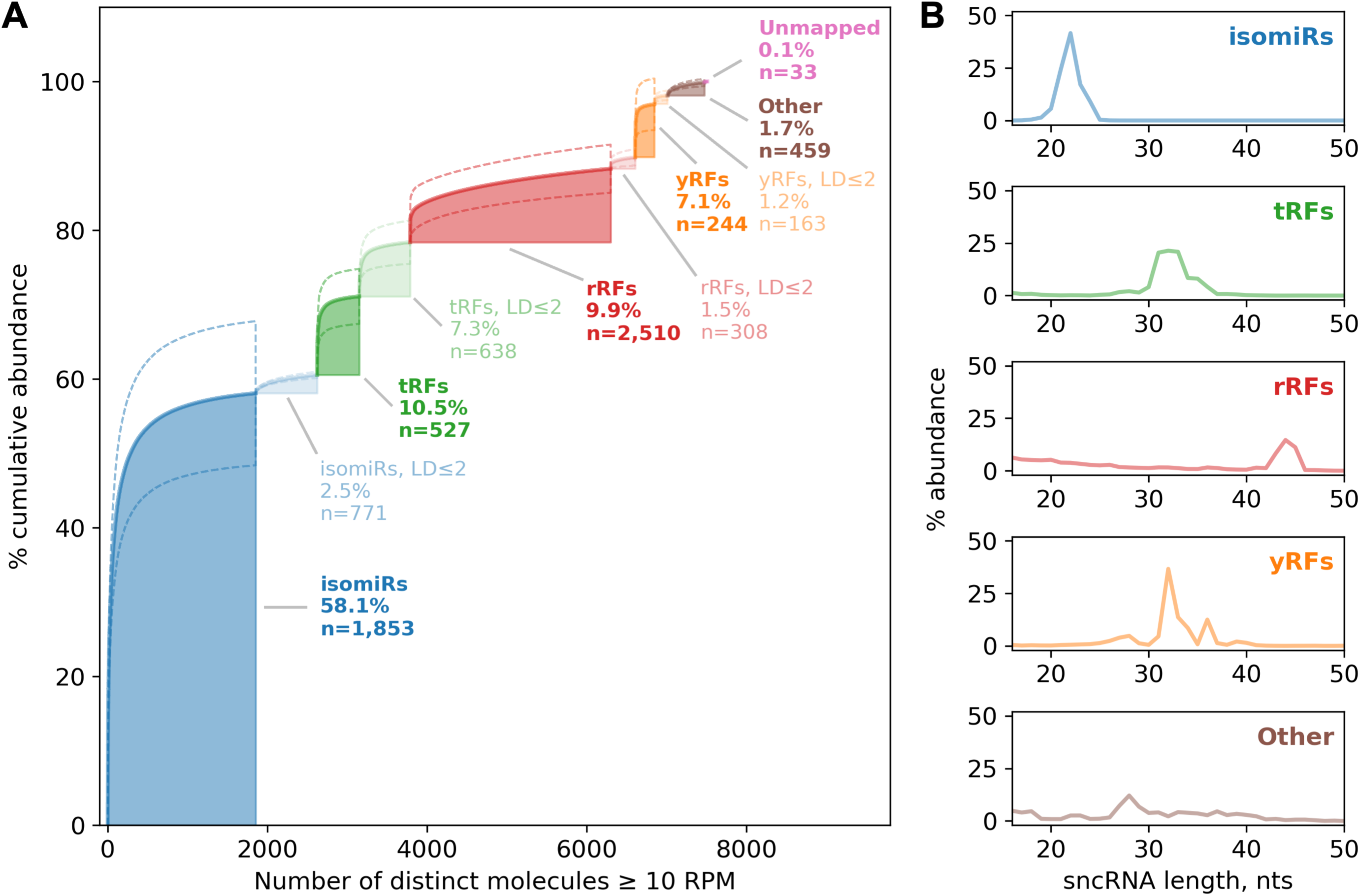
The abundance of sncRNAs in brain samples. (A) Within each molecule type, sncRNAs are ranked by mean RPM across 170 samples (rank is shown on the X-axis), and cumulative mean (solid line) ± standard deviation (dashed line) is shown on the Y-axis. “Other” molecules include rpFs and additional genomic mappings (see Methods). (B) Histograms of sncRNA lengths for each molecule type (% values sum up to 100% within each molecule type). IsomiRs, tRFs, rRFs, and yRFs include both wild-type and LD ≤ 2 molecules.

As Figure 1A shows, by profiling isomiRs, tRFs, rRFs, yRFs, and rpFs, we can annotate 99.9% of all sncRNAs whose abundance is ≥ 10 RPM. IsomiRs are the most prevalent sncRNA type, capturing 60.6% of the sncRNA-ome. The isomiRs are followed by tRFs (17.8%), rRFs (11.4%), yRFs (8.3%), and other sncRNAs (1.7%). Figure 1B shows typical lengths for each molecule type. As expected, most isomiRs are centered around a single peak of 22 nts. Most abundant tRFs have a length of 30-36 nts, suggesting that many are tRNA halves [57]. yRFs follow a similar length pattern to tRFs, whereas rRFs fall into two groups, 16-25 and 43-45 nts. Other sncRNAs (rpFs and the remaining genomic mappings) follow a “flatter” length distribution.

### Non-canonical and non-templated isomiRs abound in the brains of cases and controls

Consistently with our previous reports [31, 58], the 5′-ends of most isomiRs are canonical, i.e., they agree with the 5′ ends of the reference sequences found in public databases. Supp. Figure S3 shows detailed summary statistics on isomiR types found in the brain samples, separately for 5p and 3p miRNA arms. Several highly abundant isomiRs are notable because they have shifted “seed” regions. Since a miRNA’s/isomiR’s seed determines the identities of its targets, it follows that these isomiRs target different genes than the reference isomiR, as we showed previously [28]. Several of these isomiRs arise from the 3p arm of miR-126 and lack the first (5′-end) nucleotide of the reference sequence. The combined mean abundance of these isomiRs is high, exceeding 500 RPM. Also mirroring our previous findings, most non-canonical 5′-end isomiRs originate from the 3p arms of the miRNA precursors (Supp. Figure S3B), their 5′-ends corresponding to the point of cleavage by the DICER enzyme [31, 58].

On the other hand, more than half of all isomiRs have non-canonical 3′ termini, with ≥ 20% having non-templated nucleotide additions (Supp. Figure S3C-D). The frequencies of non-templated 3′-end nucleotides acutely differ between 5p and 3p miRNA arms, with adenylation being most frequent among isomiRs from the 5p arm (Supp. Figure S3E) and uridylation among isomiRs from the 3p arm (Supp. Figure S3F). Surprisingly, the percentage of isomiRs whose 3′-end is guanylated is significantly higher in SCZ cases (multivariable regression p-value < 3.3e-05, Supp. Figure S3E-F). Below, we will revisit this observation in more detail.

### Many tRFs with sequence mismatches abound in the brains of cases and controls

Figure 1A shows an unexpectedly high abundance of molecules with LD ≤ 2 from known tRFs (7.3% of total sncRNA-ome). In fact, this number is comparable with the abundance of known tRFs (10.5%). For simplicity of presentation, we analyzed in detail molecules with LD = 1, which constitute 84% of the LD ≤ 2 category total abundance (Supp. Table S2).

Based on the types and locations of mismatches when compared to the wild-type (WT) tRFs, we classified LD = 1 tRFs into two categories that cover 90% of all LD = 1 reads. The first category (48% of all LD = 1 reads) comprises tRFs with a non-templated nucleotide attached to the 3′-end, mirroring the concept of non-templated isomiRs. These non-templated additions are preferentially added to the 5′-halves and 5′-tRFs (90% of all tRFs with 3′-end additions) and preferentially found in tRFs from tRNA^GlyCCC/GlyGCC^, tRNA^GluCTC^, tRNA^LysCTT^, and tRNA^HisGTG^ (92% of all tRFs with 3′-end additions). Supp. Figure S4A shows the frequencies of the non-templated nucleotides. Strikingly, the distribution closely resembled the distribution for 3p miRNA arms (Supp. Figure S3F), including the statistically significant elevation of guanylation rates in SCZ cases (p-value = 1.9e-07), suggesting common processing mechanisms.

The second category is composed of extremely abundant 5′-halves and 5′-tRFs (some of them are > 1,000 RPM) from tRNA^GluCTC^ and tRNA^GlyCCC/GlyGCC^ with deletions or substitutions at position 6 of the parental tRNA (Supp. Figure S4B). We do not know whether these molecules are real or the two tRNAs have uncharacterized chemical modifications at position 6 causing incorrect sequencing.

### The sncRNA-omes of SCZ and BD cases differ significantly from controls

We begin the sncRNA-ome comparisons of cases and controls from the HBCC brain bank, as it contains both SCZ and BD cases and spans a wide age range. Using a multivariable DESeq2 model adjusted for known and hidden confounding variables (see Methods), we found 1,127 sncRNAs that differ significantly (FDR < 0.05) in abundance between SCZ cases and controls (Figure 2A-E, Supp. Table S3A). Consistently with the previous mRNA-level reports, most sncRNAs have modest FC values: median |log_2_ FC| across 1,127 DA sncRNAs is only 0.58, which is consistent with the polygenic inheritance of SCZ [59]. Considering an additional |log_2_ FC| threshold of 0.4 (filled markers in Figure 2A-E), the number of DA sncRNAs is 761.

**Figure 2.**
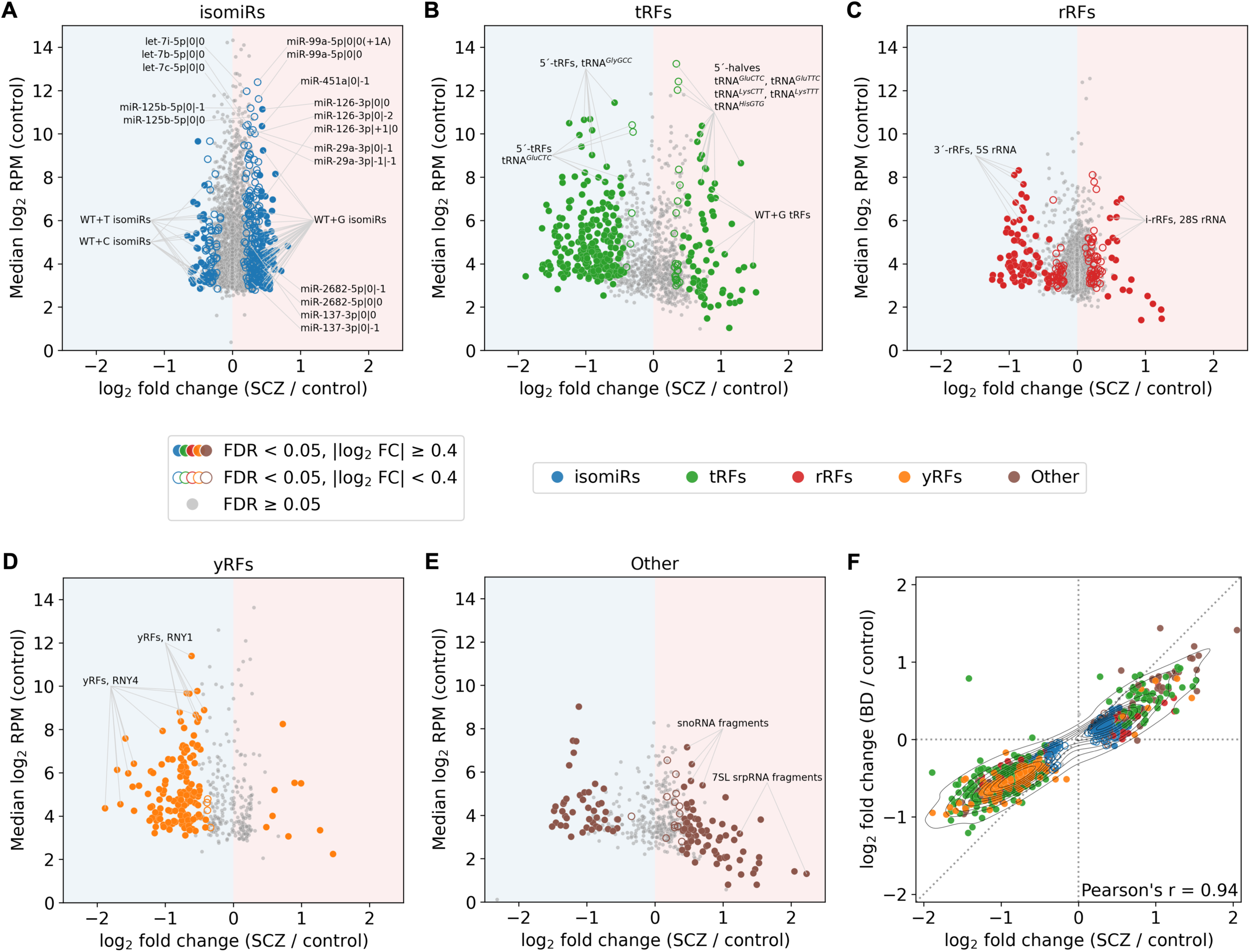
Differential abundance of sncRNAs between SCZ cases and controls in the HBCC brain bank. (A-E) Volcano plots for different sncRNA types. Significantly differentially abundant sncRNAs (FDR < 0.05) are highlighted with color and are filled if |log_2_ FC| ≥ 0.4. (F) Mutual distribution of FCs from SCZ vs. control and BD vs. control comparisons. The diagonal dotted line corresponds to equal FCs (Y = X). Only molecules associated with SCZ or BD (likelihood-ratio test FDR < 0.05) are shown. Markers are colored and/or filled if the same FDR and FC thresholding criteria are met in either of the two comparisons. IsomiRs, tRFs, rRFs, and yRFs include both wild-type and LD ≤ 2 molecules. “Other” molecules include rpFs and additional genomic mappings (see Methods).

We found a weaker effect in BD: there are only 104 DA sncRNAs between BD and controls, with all FDR values ranging between 0.01 and 0.05 (Supp. Table S3B). We further ran DESeq2 with the likelihood-ratio test to find sncRNAs that are significantly associated either with SCZ or BD (Supp. Table S3C). For these sncRNAs, the FCs between SCZ and controls are highly correlated with the FCs in the BD vs. control comparison (Pearson’s r = 0.94, p-value < 1e-100, Figure 2F). At the same time, the absolute values of FCs in the BD case are systematically lower compared to SCZ (Figure 2F).

### Highly abundant miRNAs increase further in abundance in SCZ

The effect of an isomiR on its target depends on the isomiR’s abundance. With that in mind, we focused on the DA isomiRs with exceptionally high abundance (median in SCZ or control samples ≥ 1,000 RPM). The top part of Figure 2A shows 12 such isomiRs – all of them are upregulated in SCZ; Supp. Figure S5A shows box plots for these isomiRs in SCZ cases and controls. Only 2 out of 12 isomiRs have log_2_ FC above 0.4: miR-126-3p|0|0 and miR-451a|0|-1. Notably, the previously mentioned sister isomiR miR-126-3p|+1|0 with non-canonical 5′ terminus (and, thus, different target genes) is also upregulated (Figure 2A).

The remaining isomiRs originate from the let-7 family (let-7b-5p, let-7c-5p, let-7i-5p), miR-99a-5p, miR-125b-5p, and miR-29a-3p. Three of the arms producing these miRNAs (miR-99a-5p, let-7c-5p, and miR-125b-5p) are clustered on chromosome 21 and are co-transcribed [60]. One of the two isomiRs from miR-29a-3p miRNA, miR-29a-3p|-1|-1, has a non-canonical 5′ terminus. These isomiRs are very abundant: thus, despite their modest FCs, a small change in their abundance could have considerable functional consequences.

### Differentially abundant tRFs, rRFs, and yRFs exhibit biases in the sign of their change

The tRFs, rRFs, and yRFs we identified show strong DA signatures, evident biases in the sign of their change (Figure 2B-E), and generally exhibit stronger FCs than isomiRs. Figure 2B shows a volcano plot with DA tRFs; a subset of highly abundant DA tRFs (≥ 1,000 RPM) is separately depicted in Supp. Figure S5B.

Despite the evident bias for decreased abundance in the SCZ samples, two clearly-defined groups of tRFs are upregulated in SCZ. The first group consists of highly abundant 5′-tRNA halves (three of them exceed 5,000 RPM in SCZ), including the above mentioned tRNA^GluCTC^ 5′-tRNA halves with a deletion at position 6. The second group includes tRFs with non-templated 3′ guanylation that resembles the isomiRs’ pattern. In complete analogy with isomiRs, sibling tRFs that are produced from the same parental tRNA have opposite FC signs. For example, a 34-nt 5′-tRNA half derived from tRNA^GluCTC^ increases in abundance in SCZ (log_2_ FC = 0.72, FDR = 0.01), whereas a 28-nt 5′-tRF from the same tRNA is less abundant in SCZ compared to control samples (log_2_ FC = -1.09, FDR = 0.0014). Similarly, Figure 2C shows that DA rRFs are typically less abundant in SCZ, with several clearly defined exceptions. Namely, a group of i-rRFs that are produced from different regions of 28S rRNA are more abundant in SCZ samples, whereas several other 28S i-rRFs are less abundant in SCZ. This couples SCZ to the differential processing of tRNAs and rRNAs into fragments, suggesting underlying events that go beyond changes to transcription rates.

Figure 2D shows that virtually all DA yRFs are depleted in SCZ. The most abundant yRFs originate from two parental Y RNA molecules: RNY1 and RNY4. Figure 2E depicts FCs and abundances of other sncRNAs that are not isomiRs, tRFs, rRFs, or yRFs. Of note, a 26-nt fragment of the 7SL signal recognition particle RNA (srpRNA) has the highest absolute FC across all sncRNAs (log_2_ FC = 2.23, FDR = 2e-05) albeit low average abundance (median across SCZ samples = 10.2 RPM).

### GWAS-identified miR-137-3p and miR-2682-5p are upregulated in SCZ but low in abundance

Several large-scale GWAS SCZ studies consistently found a strong, statistically significant group of SNPs proximal to miRNAs miR-137-3p and miR-2682-5p [3, 61]. Two isomiRs of miR-137-3p increase in SCZ (Figure 2A) but the effect size is limited: miR-137-3p|0|-1 (FDR = 0.041, log_2_ FC = 0.29) and miR-137-3p|0|0 (FDR = 0.082, log_2_ FC = 0.27). On the other hand, the much less studied miR-2682-5p increases statistically significantly and shows a greater magnitude of change (FDR = 0.0061, log_2_ FC = 0.45). Even when combining all DA isomiRs of miR-137-3p and miR-2682-5p, the resulting abundance is less than 50 RPM in each case. The low abundance levels and the small effect sizes of these two miRNAs suggest that their changes in SCZ are of little consequence in regard to the silencing of gene targets.

### Non-templated isomiRs differ between SCZ cases and controls

By analyzing sets of DA isomiRs, we observed clear biases in the identity of the nucleotides that are post-transcriptionally added to the 3′ end of the non-templated isomiRs (Figure 2A). Figure 3A shows the results of a GSEA analysis that assesses the enrichment of different non-templated nucleotides among differentially expressed sncRNAs. The plot’s X-axis shows the ranked list of all sncRNAs at RPM ≥ 10. The top left panel shows running enrichment scores corresponding to different non-templated isomiRs. The middle left panel shows the type of the non-templated isomiR using tick marks at the corresponding positions of the ranked list. SncRNAs that are upregulated in SCZ are on the left side, and the most downregulated ones are on the right side (bottom left panel). IsomiRs with non-templated 3′ runs of cytosine (referred to as “WT+C” isomiRs, p-value = 8e-14) and uridine (referred to as “WT+T” isomiRs, p-value = 2e-03), respectively, are prevalent among the isomiRs whose abundance *decreases* in SCZ. In fact, 80% of the *downregulated* isomiRs with log_2_ FC ≤ -0.4 belong to these two groups. IsomiRs with non-templated 3′ runs of guanine – we will refer to them as “WT+G” isomiRs – are particularly notable: they are prevalent among the isomiRs whose abundance *increases* in SCZ (p-value = 9e-27). In fact, 54% of all *upregulated* isomiRs with log_2_ FC ≥ 0.4 belong to the WT+G group. An additional 24% of isomiRs end with a guanine, but it cannot be distinguished whether the guanine is the one present on the genomic template or a post-transcriptional addition to a shorter-by-one-nucleotide templated isomiR.

**Figure 3.**
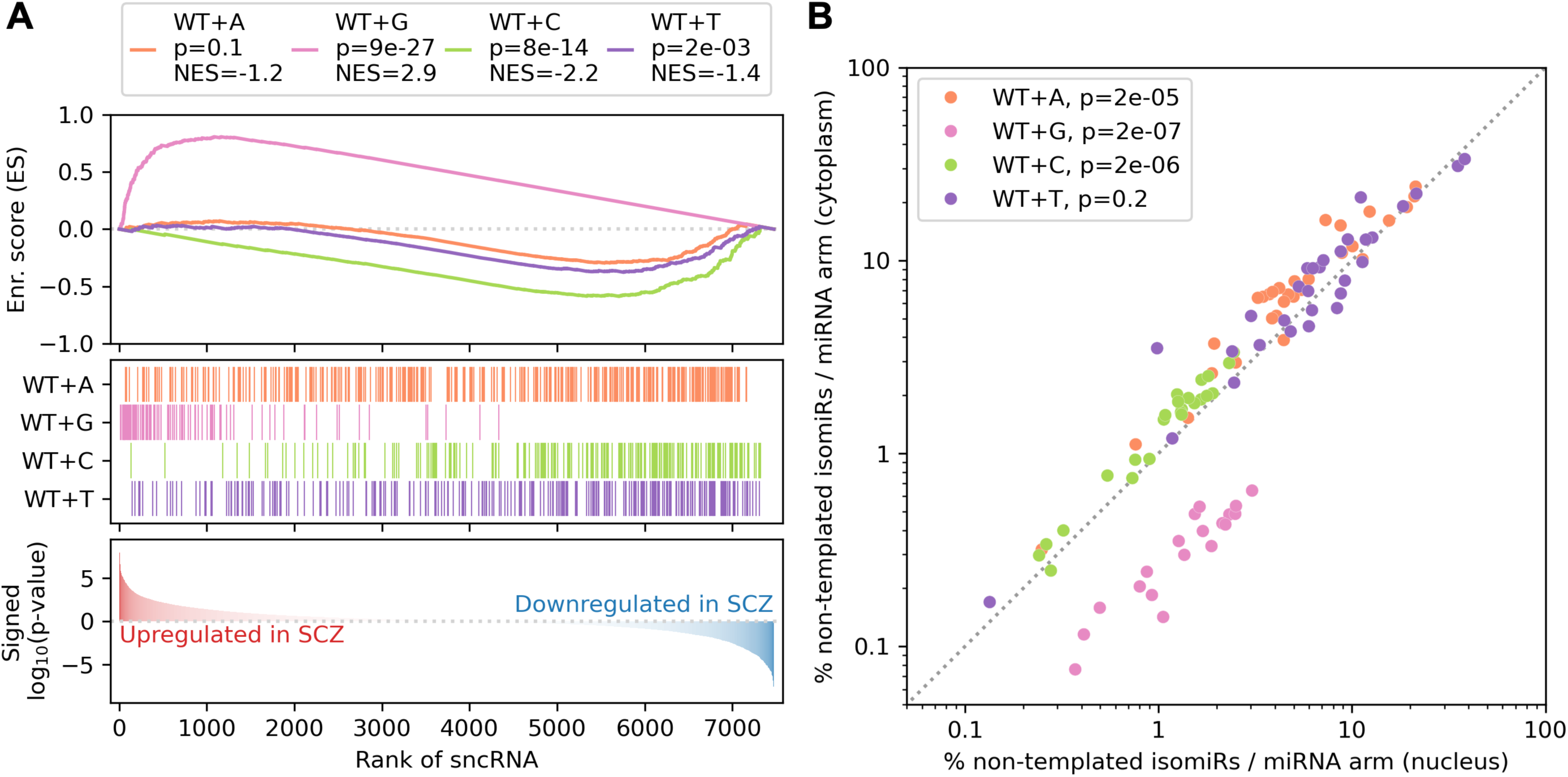
Enrichment analysis of non-templated isomiR nucleotide additions in SCZ. (A) GSEA plot for SCZ vs. control comparison in the HBCC brain bank. All abundant sncRNAs are ranked according to the log_10_ p-value multiplied by the fold change sign. Vertical bars on the middle panel indicate the presence of an isomiR with non-templated nucleotide addition. (B) Comparison of nuclear and cytoplasmic abundances of non-templated isomiRs in rat primary cortical neurons [62]. Non-templated isomiRs of 35 selected miRNA arms are shown (see main text for selection criteria). The X and Y axes show the ratio of the number of reads mapped to particular non-templated isomiR and the whole miRNA arm. The dotted diagonal line corresponds to equal FCs (Y = X).

We also found several instances where the FC of two sister isomiRs from the same miRNA arm increases or decreases depending on the identity of their non-templated nucleotide addition. For example, miR-181a-5p|0|-1(+1G) increases in SCZ compared to controls (FDR = 0.0095, log_2_ FC = 0.46), whereas miR-181a-5p|0|-2(+1C) goes down in SCZ (FDR = 0.036, log_2_ FC = -0.42). This example, together with Figure 3A, suggests that the observed DA of isomiRs can also result from differential processing of miRNAs, in addition to differential transcription.

WT+G isomiRs in the neurological context were originally reported in a previous study that deep-sequenced sncRNAs from nuclear and cytoplasmic fractions of rat cortical neurons and showed that they are enriched in the nuclear fractions [62]. Given our findings (Figure 3A), we re-analyzed the data of that study from the vantage point of isomiRs. Out of 55 miRNA arms that give rise to WT+G isomiRs with an abundance ≥ 10 RPM, 37 have identical sequences between humans and rats. 35 (94.6%) of these miRNA arms are expressed in rat neurons at an abundance of 10 RPM or higher (Supp. Table S4). Figure 3B shows the abundances of the non-templated rat isomiRs originating from these 35 miRNAs in the nuclear and cytoplasmic fractions. The WT+G rat isomiRs are highly enriched in the nuclear fraction (p-value = 2e-07), whereas the WT+C (p-value = 2e-06) and WT+A (p-value = 2e-05) isomiRs are enriched in the cytoplasmic fraction.

### The changes of mRNAs in SCZ and BD mirror the changes of sncRNAs

We also analyzed the mRNA sequencing data from the same samples that were previously reported [2]. We considered only the 15,414 mRNA genes whose abundance is at least 1 TPM in at least 25% of all (case and control) samples. There are 1,442 DA genes between SCZ and control, with only 155 genes having |log_2_ FC| ≥ 0.4 (Supp. Figure S6A, Supp. Table S5A). As in the case of sncRNAs, fold changes of SCZ- and BD-associated differences are highly correlated (Pearson’s r = 0.9, p-value < 1e-100), with the effects in BD being weaker on average (Supp. Figure S6B, Supp. Table S5B-C). To compare the changes with previously reported mRNA-level data, we applied GSEA with the reference list of DA genes generated by the PsychENCODE Consortium [1]. The analysis shows clear, highly statistically significant (p-value < 1e-200) enrichment with matching FC signs between our SCZ analysis and reference PsychENCODE SCZ data (Supp. Figure S6C). A similar highly significant concordance (p-value < 1e-60) exists between our BD FCs and PsychENCODE BD data. Gene Ontology (GO) term enrichment analysis with GSEA shows enrichment of pathways related to synaptic signaling, neuron and axon development among downregulated genes in SCZ, and translation-related pathways among upregulated genes (Supp. Figure S6D-E, Supp. Table S6).

### The basic trends are replicated in the MSSM brain bank but with weaker effect sizes

We independently ran DA tests on sncRNAs and mRNAs of the MSSM brain bank. We found only five sncRNAs (Supp. Table S3D) and one mRNA (Supp. Table S5D) whose FDR satisfied the 0.05 threshold. To compare the sign of molecule changes between the two brain banks, we used GSEA on the MSSM-ranked sncRNAs list using statistically significant HBCC changes as a reference list (Supp. Figure S7A). Both brain banks show statistically significant concordance (p-value = 2e-11), and their FCs signs generally agree (Supp. Figure S7A). All the trends with non-templated isomiRs also replicate at the level of GSEA, with WT+G isomiRs showing significantly increased abundance in SCZ and WT+A/WT+C/WT+T isomiRs showing significantly decreased abundance (Supp. Figure S7B). GSEA also shows statistically significant agreement between DA mRNA-seq results of MSSM and HBCC (Supp. Figure S7C) and between MSSM and PsychENCODE (Supp. Figure S7D). At the same time, enrichments for the PsychENCODE reference list are lower in the MSSM (Supp. Figure S7D) compared to the HBCC (Supp. Figure S6C). In summary, the MSSM brain bank replicates the HBCC-derived results but with weaker FCs.

### Both sncRNA and mRNA expression profiles suggest accelerated aging in SCZ and BD

The differences in the strength of the DA signal between the HBCC and MSSM brain banks can be due to either or both of two reasons: differences in the age distribution (MSSM individuals are significantly older) or systematic batch effects (e.g., differences in sample collection and processing). Previous studies support the first option, showing that the mRNA DA signal between SCZ cases and controls vanishes in older people [63–65].

To test the interaction between disease status and age, we conducted separate DA analyses in groups of age < 45 years old and age ≥ 45 of the HBCC brain bank. We chose the cutoff point to balance the number of cases and controls in the age groups). We also used randomized down-sampling (see Methods) to ensure that sample size does not affect the analysis (see Methods).

In concordance with the previous studies, the younger group (age < 45) is responsible for the SCZ DA mRNA signal, whereas the older group (age ≥ 45) shows a total absence of DA mRNAs (Figure 4A, “horizontal” comparisons). Interestingly, hundreds of mRNAs are DA between younger and older controls, while there are almost no differences between younger and older SCZ cases (Figure 4A, “vertical” comparisons). The same observation holds for the BD cases (Supp Figure S8A).

**Figure 4.**
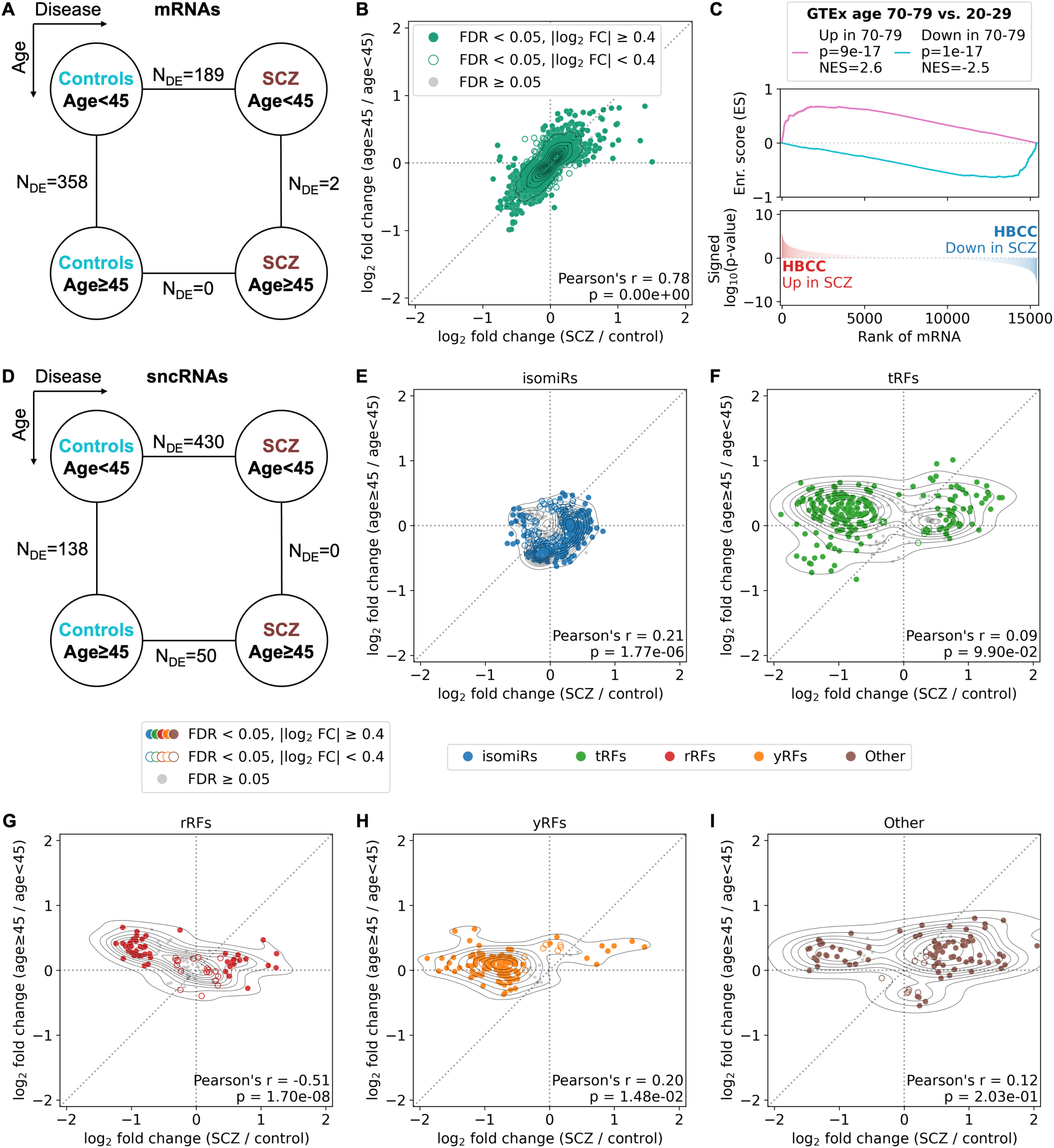
Age-dependent differential mRNA and sncRNA abundance between SCZ cases and controls (HBCC brain bank). (A, D) The number of significantly differentially abundant mRNAs/sncRNAs (FDR < 0.05) between age-restricted SCZ vs. control comparisons (horizontal dimension) and condition-restricted age ≥ 45 vs. age < 45 comparisons (vertical dimension). To remove the effect of sample size on the reported numbers, we show median numbers of differentially abundant molecules derived from 100 random down-samplings (see Methods). (B, E-I) Mutual distribution of FCs from SCZ vs. control and age ≥ 45 vs. age < 45 comparisons. The diagonal dotted line corresponds to equal FCs (Y = X). Only molecules associated with disease status or age group (likelihood-ratio test FDR < 0.05) are shown. Significantly differentially abundant mRNAs/sncRNAs are highlighted with color if FDR < 0.05 in either of the two comparisons. Highlighted markers are filled if |log_2_ FC| ≥ 0.4 in either of the two comparisons. IsomiRs, tRFs, rRFs, and yRFs include both wild-type and LD ≤ 2 molecules. “Other” molecules include rpFs and additional genomic mappings (see Methods). (C) GSEA of differentially abundant mRNAs in SCZ in the HBCC brain bank with the GTEx aging signature as a reference gene set.

Figure 4B shows the joint distribution of FCs from two contrasts of the same multivariable models: SCZ cases versus controls on the X-axis and age ≥ 45 versus age < 45 individuals on the Y-axis. Only mRNAs significantly associated with SCZ or age (likelihood-ratio test) are included (Supp. Table S5E-F). These two FCs show a striking positive correlation of 0.78 (p-value < 1e-100), suggesting protein-coding gene changes that are common to SCZ and natural aging. We observed the same striking correlation when conducting GSEA on the HBCC SCZ changes with a brain aging gene signature derived by a different group from the independent GTEx project [66] (p-value < 1e-16, Figure 4C). These observations explain the effects from Figure 4A and the weak effect sizes in the MSSM samples. Specifically, the effect of SCZ on the brain samples resembles the effect of natural aging, which minimizes the differences between SCZ and controls in older individuals. The high correlation between aging and disease FCs is also present when analyzing BD cases (Supp. Figure S8B-C).

A more complex picture exists at the sncRNA level. Figure 4D shows the numbers of significant DA sncRNAs corresponding to the interaction term of SCZ status and age group. The sncRNA data lead to the same conclusions as the mRNA diagram of Figure 4A. However, the correlations between SCZ and age group FCs are weaker and follow more complex, molecule-type-specific patterns (Figure 4E-I, Supp. Table S3E-F). For example, there is a modest, statistically significant positive FC-FC correlation of isomiR (Figure 4E) and yRF levels (Figure 4H), whereas the FC-FC correlation for rRFs is negative (Figure 4G). Supp. Figure S8D-I is the BD counterpart of Figure 4D-I, supporting very similar conclusions (the only exception is Supp. Figure S8D, showing almost no DA sncRNAs between BD and controls in the younger age group).

### Differentially abundant sncRNAs and mRNAs form distinct co-expression modules

To gain insight into the interrelations of DA molecules, we computed Pearson correlations between sncRNAs and mRNAs in an all-vs-all fashion (correlations were computed across the union of HBCC case and control samples). To ensure that correlations are not driven by major “third variables,” we explicitly residualized known factors (disease status, sex, age, RIN) and hidden confounders represented by principal components from the expression data (see Methods).

Figure 5 shows correlation heatmaps with hierarchical clustering of sncRNAs (Figure 5A) and mRNAs (Figure 5B). Even though we explicitly regressed out the disease status from the expression values, and despite using an unsupervised clustering algorithm, the clusters agree exceptionally well with the SCZ FC signs. Moreover, clustering using only control samples results in very similar modules (Rand index = 0.9 for sncRNAs and 0.84 for sncRNAs). These observations mean that DA sncRNAs and mRNAs form clusters of “entangled” molecules, which in turn suggest shared regulation and orchestrated changes in disease. Supp. Table S7 lists sncRNAs and mRNAs comprising each cluster, and Supp. Table S8 lists pathway enrichments associated with each mRNA cluster.

**Figure 5.**
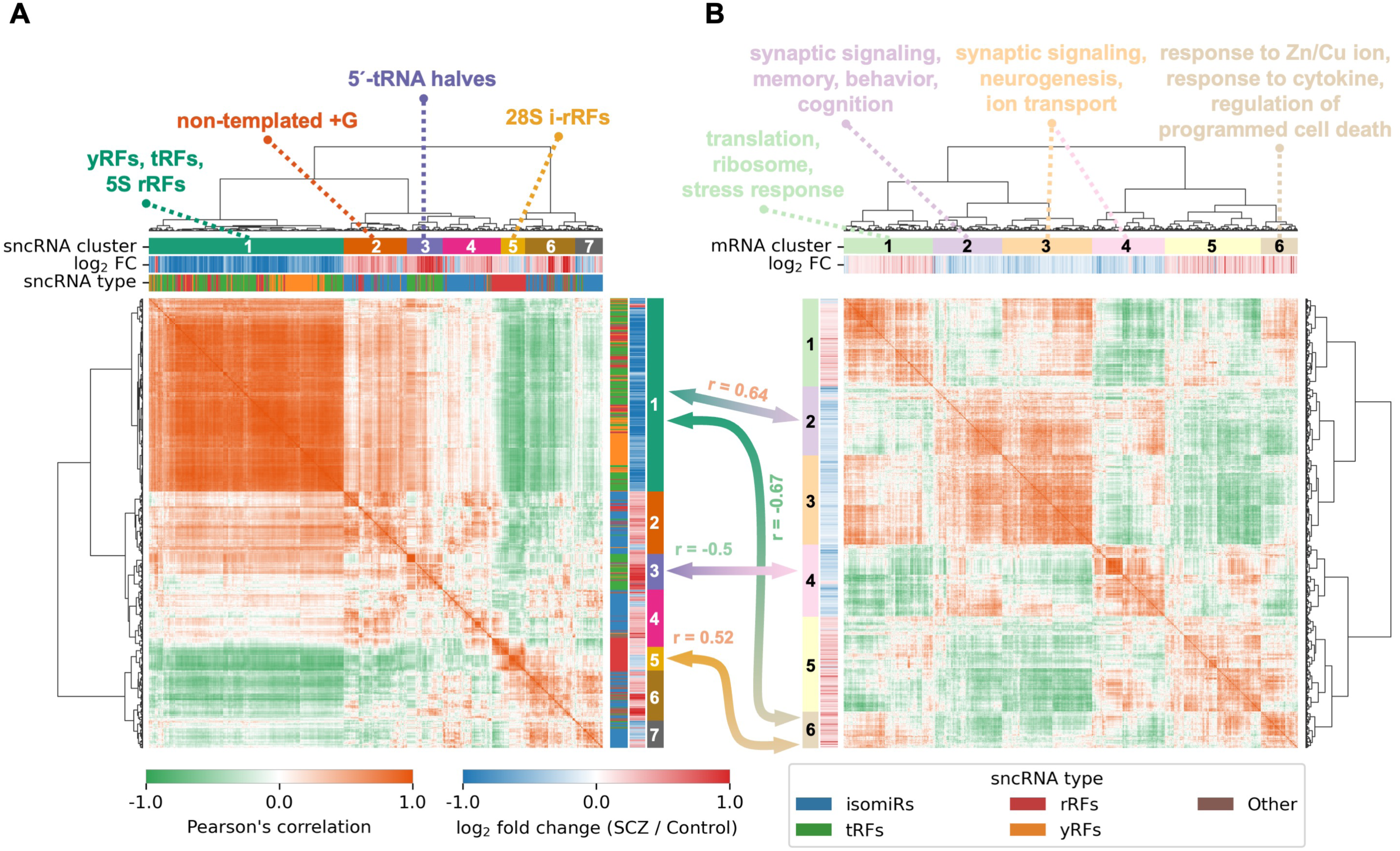
Co-expression analysis of differentially abundant sncRNAs (A) and mRNAs (B) between SCZ cases and controls in the HBCC brain bank. Correlations are computed over all HBCC samples with known and hidden confounders regressed out from the abundance levels (see Methods). Only significantly differentially abundant sncRNAs/mRNAs are included (FDR < 0.05). Annotations of mRNA clusters are based on Gene Ontology enrichment analysis. Hierarchical clustering is conducted with Ward’s linkage. Arrows between sncRNA and mRNA clusters show correlations between eigen-molecules below -0.5 or above 0.5. IsomiRs, tRFs, rRFs, and yRFs include both wild-type and LD ≤ 2 molecules. “Other” molecules include rpFs and additional genomic mappings (see Methods).

Most sncRNA clusters comprise characteristic molecule types (Figure 5A). The first cluster includes most sncRNAs whose abundance is lower in SCZ, i.e., virtually all yRFs, multiple tRFs, and rRFs predominantly derived from 5S rRNA. The second cluster is enriched in the non-templated WT+G isomiRs and is rather unexpected, as these isomiRs are grouped together by their non-templated nucleotide identity but not miRNA identity or genomic locus of origin. Cluster 3 comprises 5′-tRNA halves, whereas cluster 4 comprises various isomiRs upregulated in SCZ. Cluster 5 comprises i-rRFs derived from 28S rRNA. Lastly, clusters 6 and 7 comprise different isomiRs with increased and decreased abundance in SCZ, respectively. The structure of clusters once again highlights an additional layer of complexity independent of transcription – sncRNA processing from the precursor transcript.

Five out of six identified mRNA clusters show clear and specific GO pathway enrichments (Figure 5B). The most intriguing findings correspond to pathways with decreased abundance in SCZ samples (clusters 2, 3, 4). These include synaptic signaling, memory, behavior, cognition, neurogenesis, and ion transport. The other two clusters consist of genes upregulated in SCZ and include translation, stress response, response to Zn and Cu ions, response to cytokines, and regulation of cell death.

### SncRNA clusters are wired with specific mRNA clusters

For each identified cluster (Figure 5), we computed an “eigen-molecule,” – the first principal component based on the expression of the cluster’s genes. To link sncRNA clusters with mRNA clusters, we then computed correlations between all possible eigen-molecules.

Only four associations between sncRNA and mRNA clusters have absolute correlation values that exceed 0.5 – they are depicted by the arrows connecting the clusters between the two panels of Figure 5. The first sncRNA cluster (yRFs, tRFs, and rRFs that are downregulated in SCZ) is positively correlated with mRNA cluster 2 (downregulated in SCZ; synaptic signaling, memory, behavior, cognition) and negatively correlated with mRNA cluster 6 (upregulated in SCZ; response to ions/cytokines). The third sncRNA cluster (5′-tRNA halves increased in SCZ) is negatively correlated with the mRNA cluster 4 (downregulated in SCZ; synaptic signaling, neurogenesis). Finally, the fifth sncRNA cluster (28S i-rRFs upregulated in SCZ) is positively correlated with the mRNA cluster 6 (upregulated in SCZ; response to ions/cytokines).

### Differential abundance of multiple isomiRs, including let-7 and miR-29 families, likely causes dysregulation of their target mRNAs in mRNA clusters 2-4

While Figure 5 shows clear wirings between sncRNA and mRNA clusters, it does not answer whether sncRNAs cause mRNA changes or vice-versa. However, it is possible to generate strong evidence of causality for isomiRs by analyzing whether their predicted targets are enriched among anti-correlated mRNA that are also DA. We used two independent target prediction sources: TarBase [54] (experimentally validated targets, mainly from high-throughput CLIP data) and RNA22 [55] (purely sequence-based predictions). Since TarBase provides data only for reference miRNAs, we predicted targets with RNA22 only for isomiRs with the same mRNA-target-determining “seed” region as the corresponding reference miRNAs.

For 790 out of 1,501 expressed isomiRs (53%), we observed a statistically significant (FDR < 0.05) overrepresentation of TarBase experimentally validated targets among the 5% of genes with the most negative correlations with the isomiR (Supp. Table S9). This includes all but one (miR-451a) DA isomiRs expressed at ≥ 1,000 RPM (Figure 2A). For RNA22 predictions, this number is lower (480 out of 1,501 isomiRs, 32%), with weaker enrichments (p-value < 1e-46). We thus focused on TarBase-derived findings (790 isomiRs and their targets) for the downstream analysis.

We next analyzed whether target mRNAs that are anticorrelated with DA isomiRs tend to change in the opposite direction in SCZ. For this analysis, we residualized the disease status from the expression values, and thus, the anticorrelation does not automatically imply the opposite polarity of changes in disease. For 45 DA isomiRs (41 upregulated and 4 downregulated in SCZ), the number of DA anti-correlated target genes that changed in the opposite direction significantly exceeds what is expected by chance (FDR < 0.05), while there is no enrichment for co-directional changes for the same isomiRs (Supp. Table S10A-B). The strongest enrichment signal pertains to 13 isomiRs from the let-7 family, including canonical let-7b-5p, let-7c-5p, and let-7i-5p miRNAs with abundance ≥ 1,000 RPM (Figure 2A). Another isomiR from the same extreme abundance group, miR-29a-3p|0|-1, also shows statistically significant enrichment together with related isomiRs from the miR-29 family.

The target genes of these isomiRs are non-uniformly distributed across the mRNA clusters 2-4 containing genes with decreased expression in SCZ (Figure 5B, Supp. Table S10C). In concordance with the mRNA enrichments shown in Figure 5B, several critical pathways are highly enriched in the downregulated target gene sets, including synaptic signaling, neurogenesis, behavior, memory, cognition, and others (Supp. Table S10D).

## DISCUSSION

We generated and analyzed the largest collection, to date, of small RNA-seq datasets from post-mortem brain samples of SCZ and BD cases and controls. Using a state-of-the-art read mapping pipeline, we annotated virtually all sncRNAs with RPM ≥ 10. More than 98% of the sncRNAs belong to one of four classes (Figure 1A): isomiRs (60.6%), tRFs (17.8%), rRFs (11.4%), and yRFs (8.3%). Each sncRNA class has its own characteristic length distribution with notable peaks around multiples of 11 nts (Figure 1B): isomiRs at 22 nts, tRFs and yRFs at 33 nts, and rRFs at 44 nts.

The detailed analysis of the most abundant sncRNA type, isomiRs, revealed that most of them are non-canonical (Supp. Figure S3A-D), mirroring our previous observations from various cell types and tissues [18, 28, 31, 32, 58]. A relatively small portion of isomiRs has non-canonical 5′-ends: because these isomiRs have shifted “seed” regions, they target different sets of genes compared to the canonical isomiR [28]. One notable miRNA with a non-canonical 5′-end isomiR is miR-126-3p. It has two sets of highly abundant isomiRs: one with the canonical 5′-end and one with the 5′-end lacking the first nucleotide. Interestingly, we recently found the same two isomiRs of miR-126-3p to be abundant in megakaryocytes and platelets [31].

We also identified many highly abundant unusual tRFs that were not reported in previous large-scale analyses [44]. Nearly half of them are tRFs with a non-templated nucleotide attached to their 3′-ends (Supp. Figure S4A) and unrelated to the well-known CCA addition to mature tRNAs. The distribution of these added nucleotides mirrors the one observed at the 3p arm of isomiRs (Supp. Figure S3F). In most cases, these additions occur to 5′-tRNA halves and 5′-tRFs. A second type of unusual tRFs is characterized by a deletion at position 6 in tRNA^GluCTC^ or by a deletion/substitution at the same position in tRNA^GlyCCC/GlyGCC^ (Supp. Figure S4B). While it is conceivable that these changes may result from sequencing errors of decorated tRNA bases, we are not aware of any *known* modifications at this position. tRFs belonging to this second type were recently reported to be DA in syncytiotrophoblast extracellular vesicles in preeclampsia [67].

When comparing SCZ cases with controls (HBCC samples), we found 1,127 sncRNAs (Figure 2A-E) and 1,442 mRNAs (Supp. Figure S6A) with statistically significant differences in abundance (FDR < 0.05). The FCs in both cases are modest, in agreement with existing studies and the polygenic genetic architecture of SCZ [1, 59]. When comparing BD cases with controls we find the same sncRNAs and mRNAs having FCs highly similar to those in SCZ: correlation between SCZ-vs.-control and BD-vs.-control FCs exceeds 0.9 for both sncRNAs and mRNAs (Figure 2F, Supp. Figure S6B). This is also concordant with existing transcriptomic and genetic studies [1, 68]. At the same time, the absolute values of FCs are, on average, lower in BD, resulting in only a few sncRNAs and mRNAs that satisfy the FDR threshold of 0.05.

In addition to the HBCC, we compared SCZ cases with controls in the independent MSSM brain bank. The two brain banks show a statistically significant correlation of FCs both for sncRNAs (Supp. Figure S7A) and mRNAs (Supp. Figure S7C). However, only a handful of molecules passed the statistical significance criteria in the MSSM samples with very limited effect sizes. To determine whether the attenuated DA signal is explained by the much higher average age of the MSSM individuals, we dichotomized the HBCC into individuals < 45 and ≥ 45 years old. Strikingly, most disease DA signal comes from the younger age group (Figure 4A, 4D). Moreover, changes in mRNAs between SCZ cases and controls are highly correlated with changes associated with aging – this is true both for aging signatures derived from our cohort (Figure 4B) and independent GTEx data (Figure 4C). However, the same correlations computed over sncRNAs are not as high, and they depend on the sncRNA type (Figure 4E-I). These observations agree with the existing model of accelerated aging induced by SCZ and BD – an effect observed with different types of biomarkers [69], including the recent analyses of brain sample coding transcriptomes at bulk [63–65] and single-nucleus [70] resolution.

We are aware of three studies involving small RNA-seq analysis of post-mortem brain samples of SCZ cases. The study by Hu et al. has the largest sample size and compared miRNA profiles between the post-mortem dorsolateral prefrontal cortex of 34 SCZ cases and 102 controls [71]. This study found only two miRNAs with elevated expression in SCZ, miR-936 and miR-3162. However, it did not report baseline abundances of these miRNAs and did not explicitly mention the use of abundance cutoffs. The study by Ragan et al. compared sncRNAs isolated from the post-mortem anterior cingulate cortex of 22 SCZ cases and 22 matched controls and found no DA molecules [72]. Finally, the study by Liu et al. compared post-mortem amygdala samples of 13 cases and 14 controls and found 17 DA miRNAs, with only miR-451a being among our DA isomiRs [73]. One possible reason that these studies found few or no DA miRNAs may be that all included older controls, on average, than our HBCC controls.

To shed light on dependencies between DA molecules, we computed correlations between expression levels of all sncRNAs and mRNAs in the HBCC brain bank. We found that both sncRNAs and mRNAs cluster into distinct co-expression modules, mainly containing molecules with the same SCZ FCs sign (Figure 5). This is notable because it emerged after our explicit residualization of disease status and other confounders from the expression data. We also observed a very similar clustering when we computed correlations using only the control samples. Thus, the differential abundance of some gene groups in SCZ could be explained by the fact that these genes are tightly co-expressed in control brains and thus change in disease together as a single module.

IsomiRs are the only sncRNA species with a well-understood mechanism of gene expression regulation [4]. We thus analyzed those mRNAs that were significantly anti-correlated with isomiRs. In most cases, we found a strong and statistically significant overrepresentation of experimentally validated miRNA targets among these genes (Supp. Table S9). We also observed another highly statistically significant trend: experimentally validated target genes that are anti-correlated with DA isomiRs tend to change in the opposite direction in SCZ (Supp. Table S10A). This observation is not a trivial implication of anti-correlation because prior to correlation computations, we eliminated the disease status from sncRNA and mRNA abundance values. These sets of DA isomiR target genes belong to critical processes, including synaptic signaling, neurogenesis, behavior, memory, cognition, and others (Supp. Table S10C-D). Taken together, this constitutes strong, albeit indirect, evidence that changes in the abundances of isomiRs in SCZ are tightly coupled to the opposite polarity changes in their target genes, the latter being involved in key brain pathways.

Among the isomiRs with the strongest links to their target genes are four very abundant (≥ 1,000 RPM) ones whose abundance increases further in SCZ: let-7b-5p, let-7c-5p, let-7i-5p, and miR-29a-3p. Despite the respective FCs being modest (see upper-right part of Figure 2A), the corresponding absolute changes are substantial because of high starting abundances. The largest miRNA microarray study of post-mortem brain samples of SCZ cases we are aware of also shows the upregulation of let-7 and miR-29 families in SCZ [8]. The let-7 family is also implicated in neurodegenerative disorders [74, 75]. Both let-7 and miR-29 family miRNAs are known to be essential regulators of nervous system development and function [76–78].

Notably, other sncRNA types, especially tRFs (Figure 2B) and yRFs (Figure 2D), have DA molecules with higher abundances and stronger effect sizes than isomiRs. Using rigorous co-expression analysis, we found several strong associations between the clusters comprising tRFs/yRFs and mRNA clusters associated with the critical nervous system and other pathways (Figure 5). With a few exceptions, the mode of action of tRFs and yRFs is largely unknown. Consequently, we could not apply computational methods to untangle the directionality of interactions between these sncRNAs and protein-coding genes as we did in the case of miRNAs. Previous studies found associations between altered tRF abundances and neurodegenerative disorders and ischemic stroke response [79].

One of the strongest SCZ-associated genomic loci is proximal to clustered miRNAs miR-137-3p and miR-2682-5p [3, 61]. Interestingly, there is a statistically significant increase in the abundance of isomiRs originating from these two miRNAs in SCZ samples. However, both miRNAs have low abundance (< 50 RPM) and weak effect sizes (FCs ≤ 0.45), suggesting little to no impact on downstream mRNA targets. Concordantly, genes anti-correlated with both miRNAs are not enriched in experimentally validated targets from TarBase. We emphasize that miRNA targeting is cell-type dependent [80] and, thus, our remarks apply only to the analyzed brain regions; these miRNAs might still have an important function in SCZ in other brain regions or cell types.

One of the most unexpected findings pertains to the rearrangement of the frequencies of non-templated isomiR nucleotides in SCZ. The strongest signal comes from post-transcriptional guanylation of the isomiRs’ 3′ end, which is strikingly enriched among sncRNAs with elevated abundance in SCZ (Figure 3A). The effect is so consistent that it can also be observed when analyzing *all* isomiRs ending in non-templated guanine across *all* miRNA arms (Supp. Figure S3E-F). Guanylated isomiRs form a separate co-expression cluster (Figure 5A), meaning that the effect of non-templated guanine addition outweighs the effect of common transcription of sister isomiRs. We illustrate this phenomenon using two isomiRs from different miRNA arms located on different chromosomes, miR-99a-5p|0|-2(+1G) and miR-26a-5p|0|-1(+1G) (Supp. Figure S9). Both isomiRs belong to the same cluster and are highly correlated (r = 0.79, p-value = 5.8e-23). At the same time, correlations of the expression levels of these isomiRs with their most abundant sister isomiRs are much lower: r = 0.58 (p-value = 1.3e-10) between miR-99a-5p|0|-1 and miR-99a-5p|0|-2(+1G) and r = 0.59 (p-value = 8.1e-11) between miR-26a-5p|0|0 and miR-26a-5p|0|-1(+1G).

Khudayberdiev et al. generated deep sncRNA sequencing data from nuclear and cytoplasmic fractions of rat cortical neurons [62]. The authors reported an overrepresentation of guanine nucleotides at the 3′ termini of nuclearly-enriched isomiRs. We reanalyzed these data by focusing on miRNAs with perfect sequence conservation between humans and rats. All but two miRNAs with WT+G isomiRs in human samples are also abundant in the rat samples, demonstrating the comparability of the two data sources. Figure 3B shows a strong, almost 10-fold enrichment of guanylated isomiRs in the nuclear fraction, suggesting that changes in SCZ might affect both abundance and cellular localization of isomiRs. Interestingly, only 22% of the WT+G isomiRs show significant enrichment of experimentally validated targets of the WT miRNA among anti-correlated mRNAs, suggesting a different mode of action for these isomiRs.

There are at least two potential biogenesis pathways for guanylated isomiRs and tRFs: non-templated nucleotide addition (also called tailing) and A-to-I RNA editing. The reports on isomiR tailing with guanine are very scarce. To the best of our knowledge, there is only one study that sheds light on potential proteins involved in that process: Yang et al. showed that TENT2 partially contributes to the guanylation of isomiRs [27]. At the same time, TENT2 also adenylates and uridylates isomiRs, which does not match the direction of change of WT+A and WT+T isomiRs we observed, suggesting the existence of other, perhaps neuron-specific, mechanisms involved in isomiR tailing. We think RNA editing is an unlikely cause for the observed WT+G isomiRs since the reads with A-to-G mismatches in the beginning/middle of isomiRs are an order of magnitude fewer, and the DA signal is observed only when the last nucleotide is affected. Also, Alon et al. reported only 19 statistically significant miRNA editing sites in the human brain [81], while we observed non-templated 3′-end guanines for virtually all highly abundant miRNAs.

Our analysis of non-templated nucleotides uncovered an interesting phenomenon: two isomiRs originating from the same miRNA precursor could behave differently in disease, in some cases having opposite FC signs. For example, guanylated isomiR miR-181a-5p|0|-1(+1G) significantly increases in abundance in SCZ, while the cytosylated miR-181a-5p|0|-2(+1C) decreases. The most commonly used approach for miRNA DA analysis, which consists of using miRNA-level read counts, would miss most of this DA signal since opposing changes would cancel out during the summation of sibling isomiRs. The same applies to microarray and standard PCR-based miRNA analyses that cannot distinguish between individual isomiRs [82]. Notably, all three aforementioned small RNA-seq studies of SCZ brain samples used this miRNA-level analysis strategy [71–73]. We and others have already reported the critical importance of isomiR-level DA in cancer, COVID-19, and other contexts [31, 32, 83, 84]. Notably, this concept extends from isomiRs to other sncRNA types: tRFs and rRFs. For example, the longer 34-nt 5′-tRNA half from tRNA^GluCTC^ is upregulated in SCZ, whereas the shorter 28-nt 5′-tRF from the same tRNA is downregulated. Concordantly, these two tRFs are not correlated (r = -0.09, p-value = 0.4) and belong to different clusters (clusters 3 and 1, respectively). We recently reported the same behavior of tRNA^GluCTC^ tRFs in colon and breast cancer tissues and cell lines [57].

## Author contributions

I.R. and P.R. designed and supervised the study. I.N., P.L., and S.N. developed the small RNA-seq mapping pipeline. Z.S, J.F.F. and G.V. participated in the selection of samples and isolation of RNA. S.N., P.L., and I.R. analyzed the data and interpreted the results with contributions from P.R. and K.G. S.N. and I.R. prepared the figures and tables. S.N. and I.R. wrote the manuscript with contributions from P.R. and K.G. All authors approved the final version of the manuscript.

## Acknowledgments

The work of the Thomas Jefferson University team was supported by University funds (I.R) and NIH grant R01HG012784 (I.R.). The work of the Mount Sinai School of Medicine team was supported by NIH grants R01MH109677, U01MH116442, R01MH110921, R01MH125246, R01MH109897, R01AG067025, R01AG065582, R01AG050986 (P.R). We also acknowledge the use of the Cancer Genomics Shared Resource at the Sidney Kimmel Cancer Center (SKCC) of Thomas Jefferson University: SKCC is supported by an NIH Cancer Center Support Grant P30CA056036. We thank Dr. Gerhard Schratt for providing us with raw small RNA sequencing data of rat neurons from the study [62].

## Data availability

Raw data (FASTQ files) and processed data (read count matrices, metadata) are available in Synapse under synID syn63862703.

## SUPPLEMENTAL FIGURE CAPTIONS

**Supp. Figure S1.**
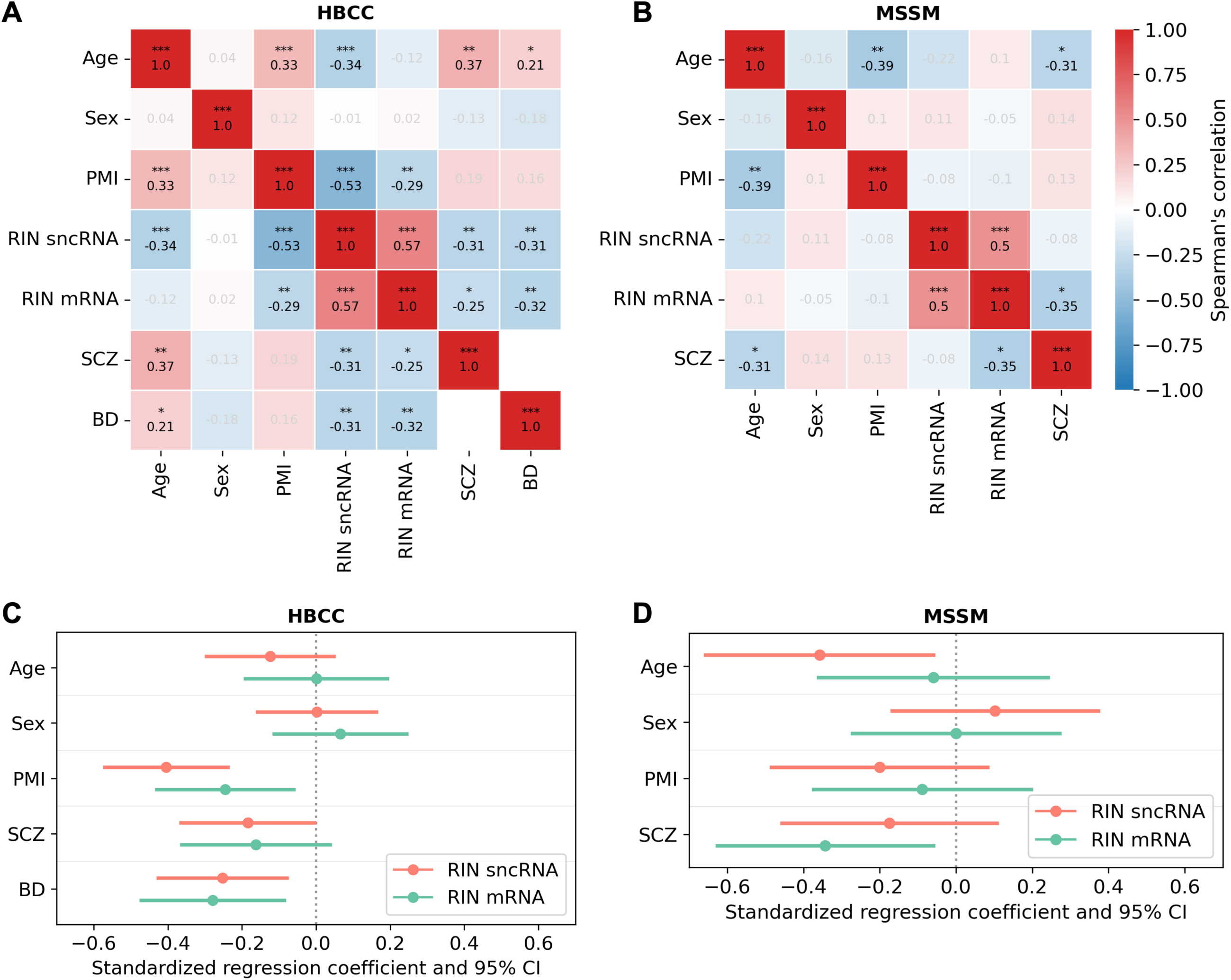
The relationships between clinical and demographical variables. (A-B) Spearman’s correlation matrices for key variables in the HBCC and MSSM brain banks. * p-value ≤ 0.05, ** p-value ≤ 0.01, *** p-value ≤ 0.001. For two binary variables (BD, SCZ, sex), p-values were computed using Fisher’s exact test. For two continuous variables (age, PMI, RIN sncRNA, RIN mRNA), p-values were computed using Spearman’s correlation test. For a binary and a continuous variable, p-values were computed using Mann-Whitney’s U-test. A positive correlation with sex means higher values among males. (C-D) Multivariable linear regression coefficients and 95% confidence intervals with RIN as a dependent variable in the HBCC and MSSM brain banks.

**Supp. Figure S2.**
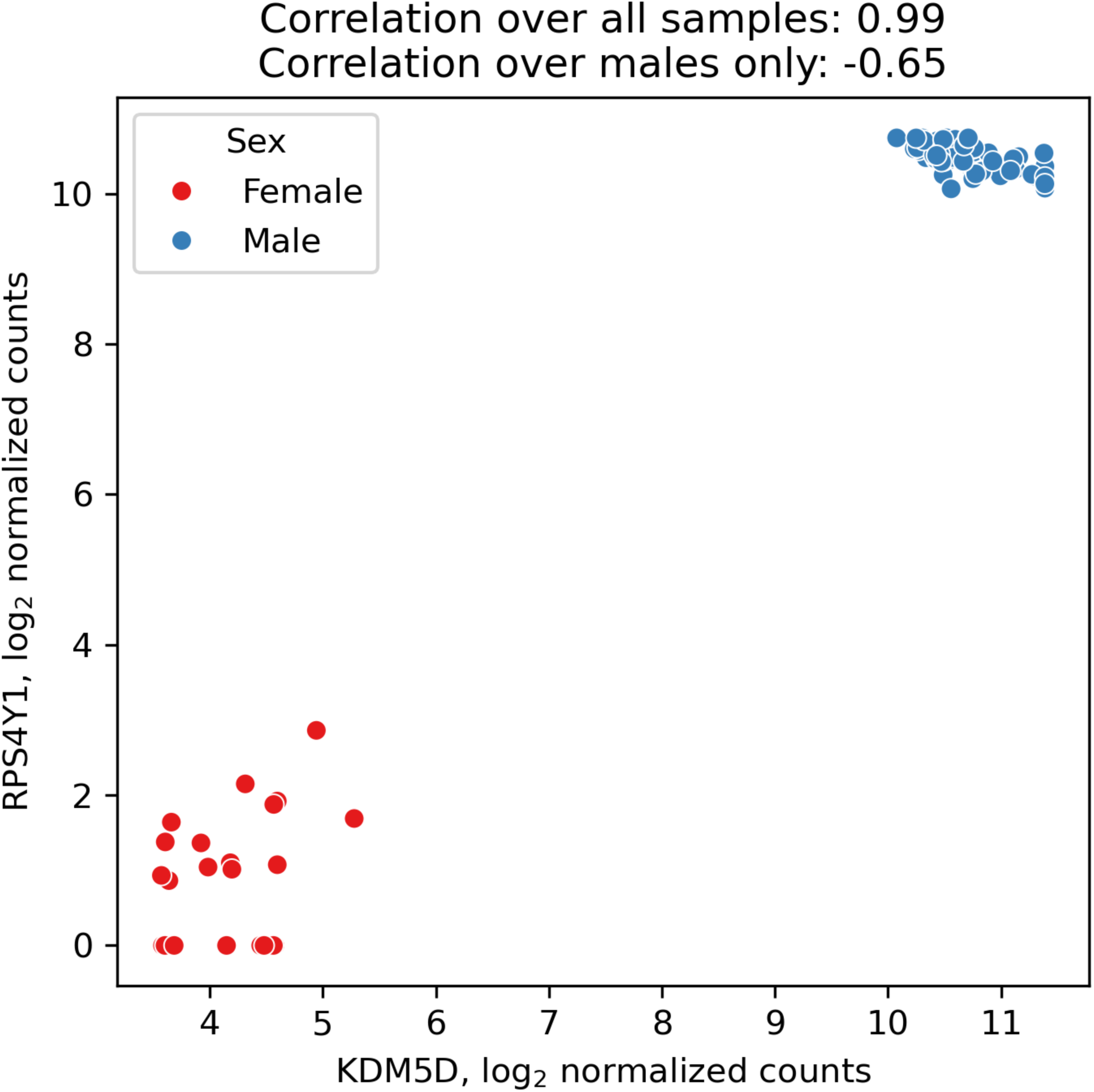
An illustration of how a confounder variable can lead to a false positive correlation. The plot shows the joint distribution of expression levels of two genes located on the Y chromosome (all samples from the HBCC brain bank). Sex variable creates a spurious positive correlation between expression levels, while the actual correlation across male samples is negative.

**Supp. Figure S3.**
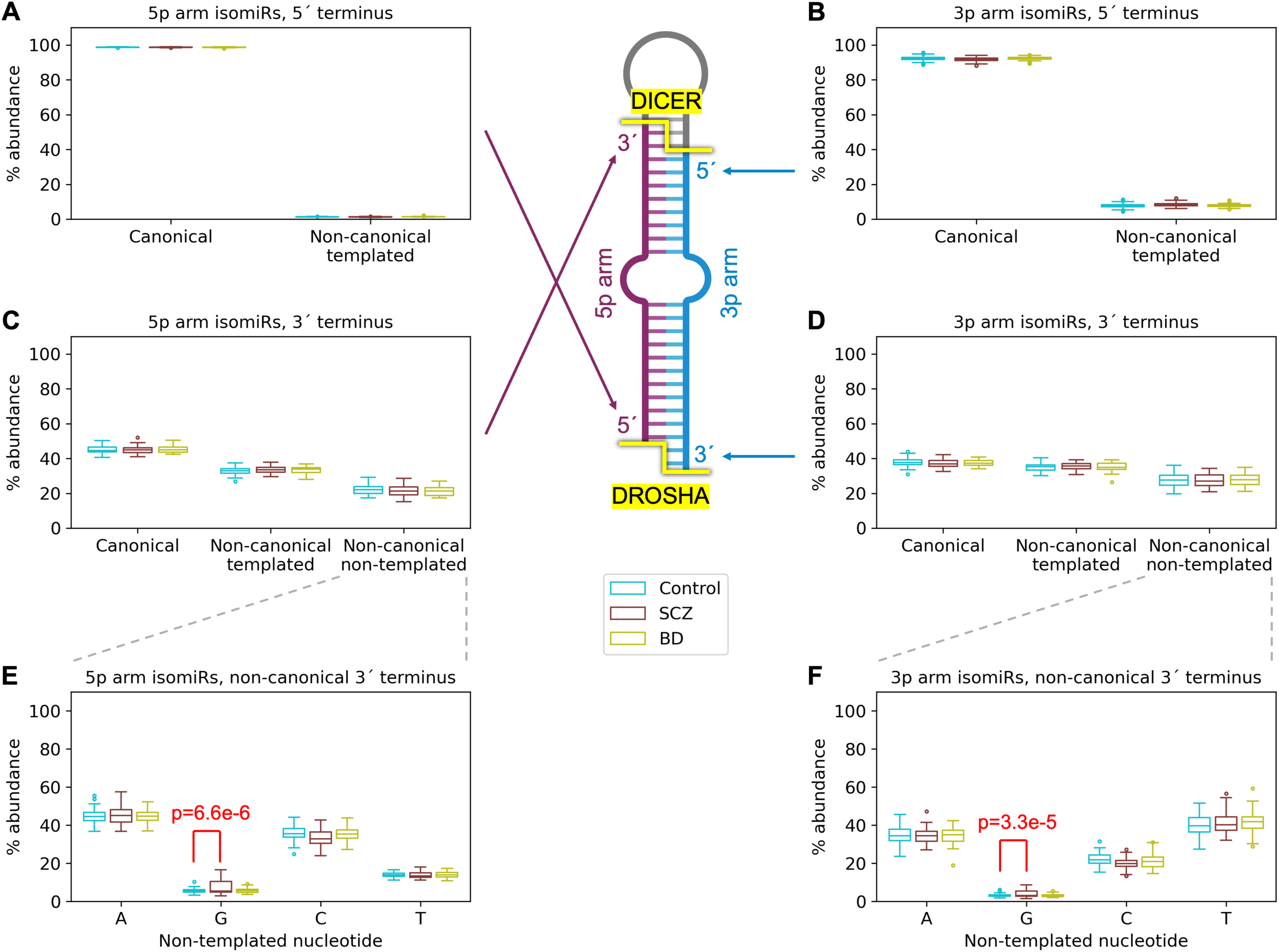
Abundances of different isomiR types. (A-D) The percentage of canonical and non-canonical isomiRs is shown separately for the four cleavage positions of precursor miRNA hairpins by DROSHA and DICER. The percentages are computed over *all* isomiRs originating from 5p miRNA arms for panels (A, C) and from 3p miRNAs for panels (B, D). (E-F) The percentage of different non-templated nucleotides at 3′-ends of isomiRs. The shown p-values are computed from a multivariable linear regression model adjusting for sex, age, RIN, and brain bank. The miRNA hairpin was created at BioRender.com.

**Supp. Figure S4.**
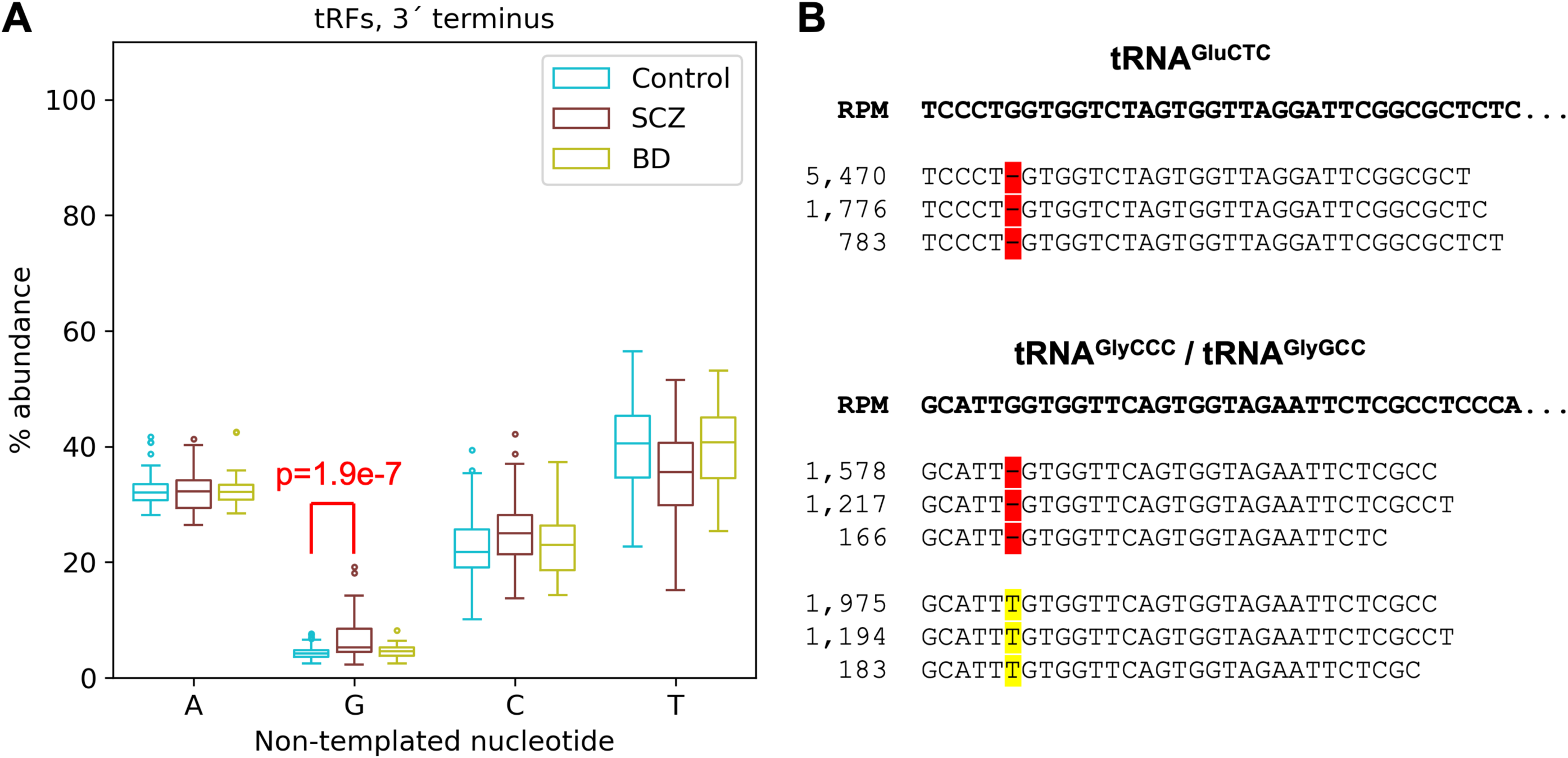
Abundances of tRFs with mismatches. (A) The percentage of different non-templated nucleotides attached to the 3′-ends of the known tRFs. The shown p-value is computed from a multivariable linear regression model adjusting for sex, age, RIN, and brain bank. (B) Highly abundant tRFs produced from tRNA^GluCTC^ and tRNA^GlyCCC/GlyGCC^ with internal mismatches at position 6. The bold nucleotides show the sequences of wild-type tRNAs. We show the 3 most abundant fragments and their mean RPMs for each type of mismatch.

**Supp. Figure S5.**
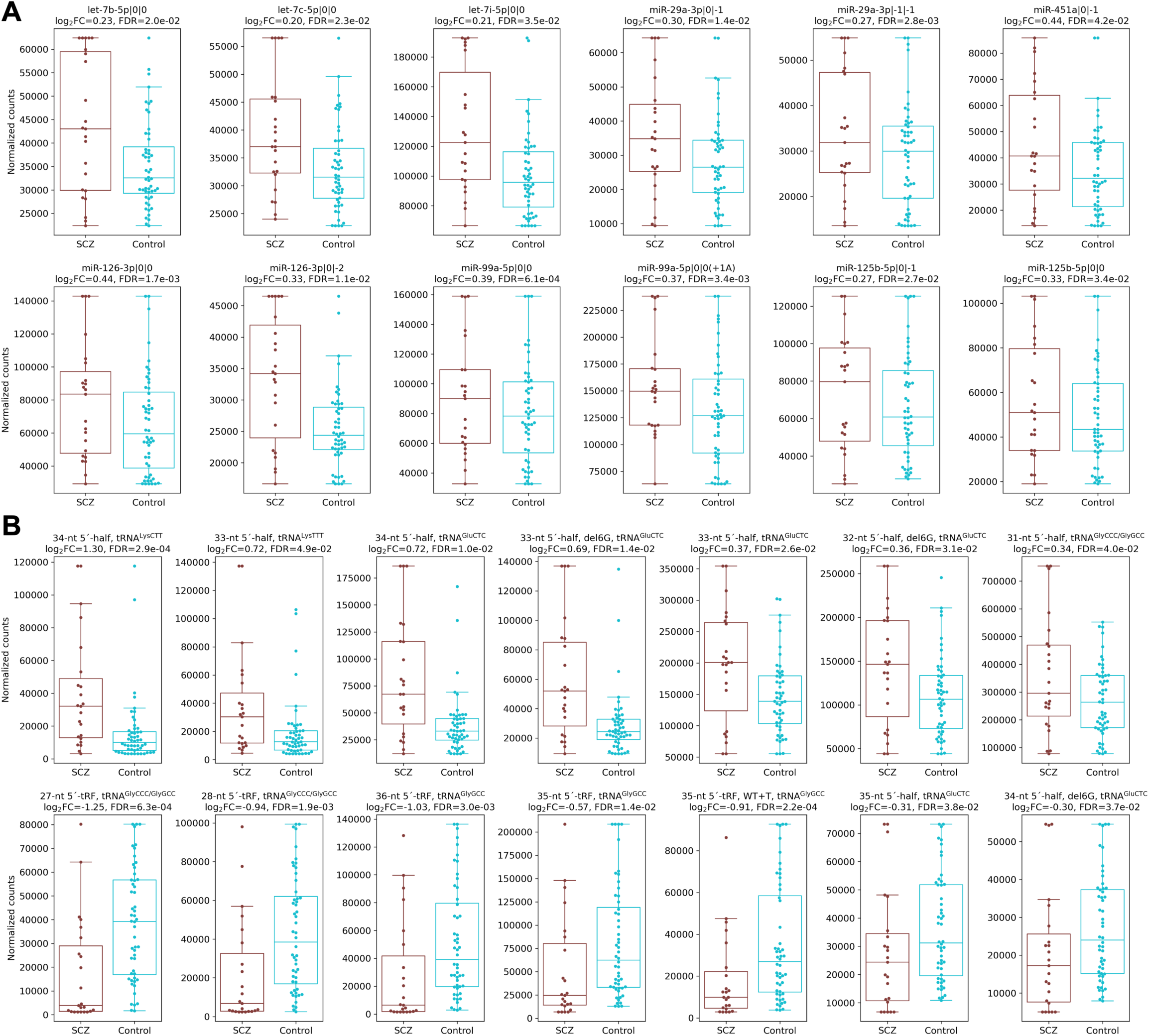
Distributions of selected DA isomiRs (A) and tRFs (B) in SCZ cases and controls in the HBCC brain bank. All molecules with FDR < 0.05 and median abundance above 1,000 RPM in either cases or controls are shown.

**Supp. Figure S6.**
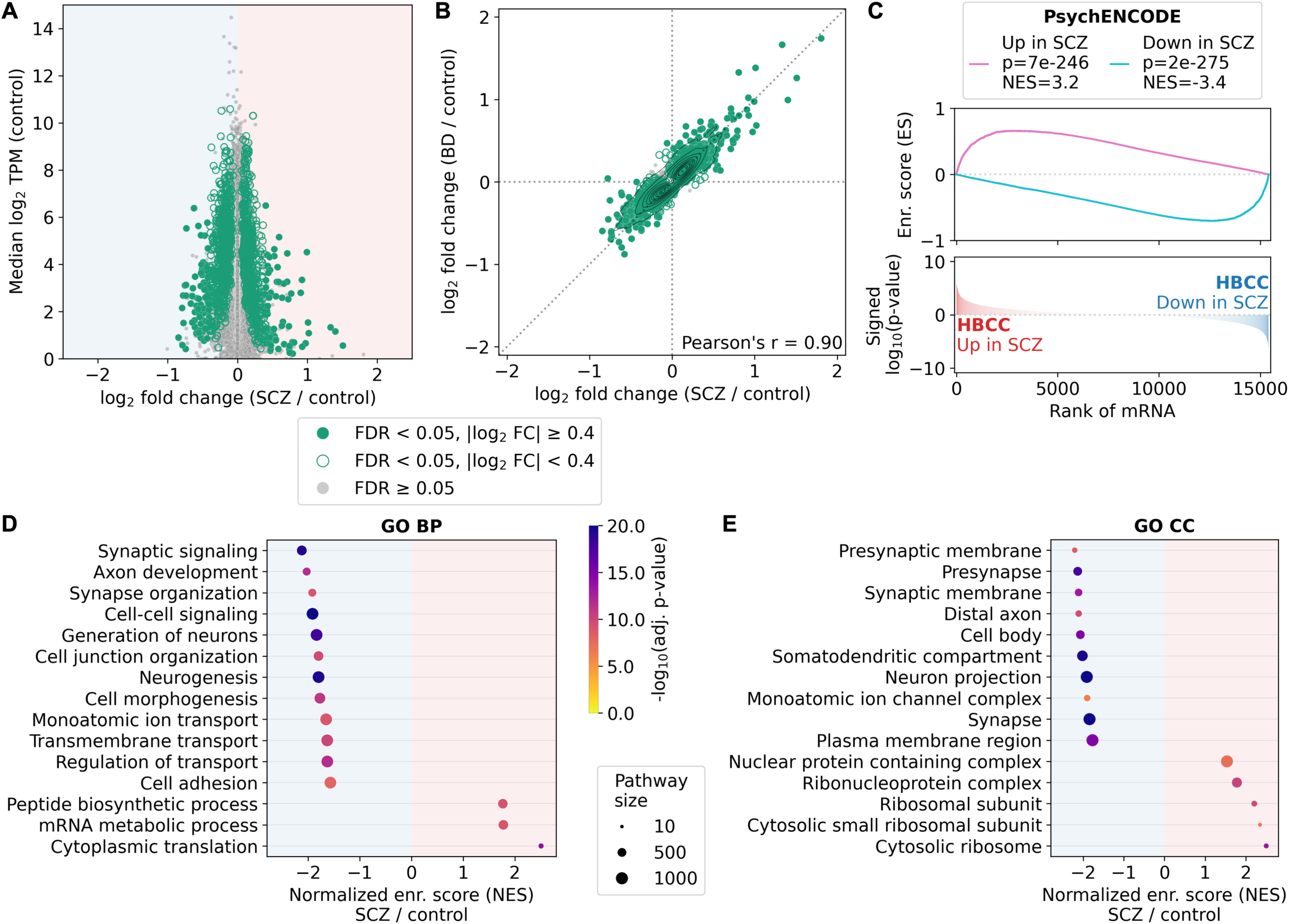
Differential abundance of mRNAs between SCZ cases and controls in the HBCC brain bank. (A) Volcano plot. Significantly differentially abundant sncRNAs (FDR < 0.05) are highlighted with green and are filled if |log_2_ FC| ≥ 0.4. (B) Mutual distribution of FCs from SCZ vs. control and BD vs. control comparisons. The diagonal dotted line corresponds to equal FCs (Y = X). Only mRNAs associated with SCZ or BD (likelihood-ratio test FDR < 0.05) are shown. Markers are colored and/or filled if the same FDR and FC thresholding criteria are met in either of the two comparisons. (C-E) GSEA with (C) the reference PsychENCODE set of differentially abundant mRNAs; (D-E) the Gene Ontology biological process (BP) and cellular component (CC) gene sets. NES stands for normalized enrichment score. Only non-redundant pathways selected by the “collapsePathways” function from fgsea are shown.

**Supp. Figure S7.**
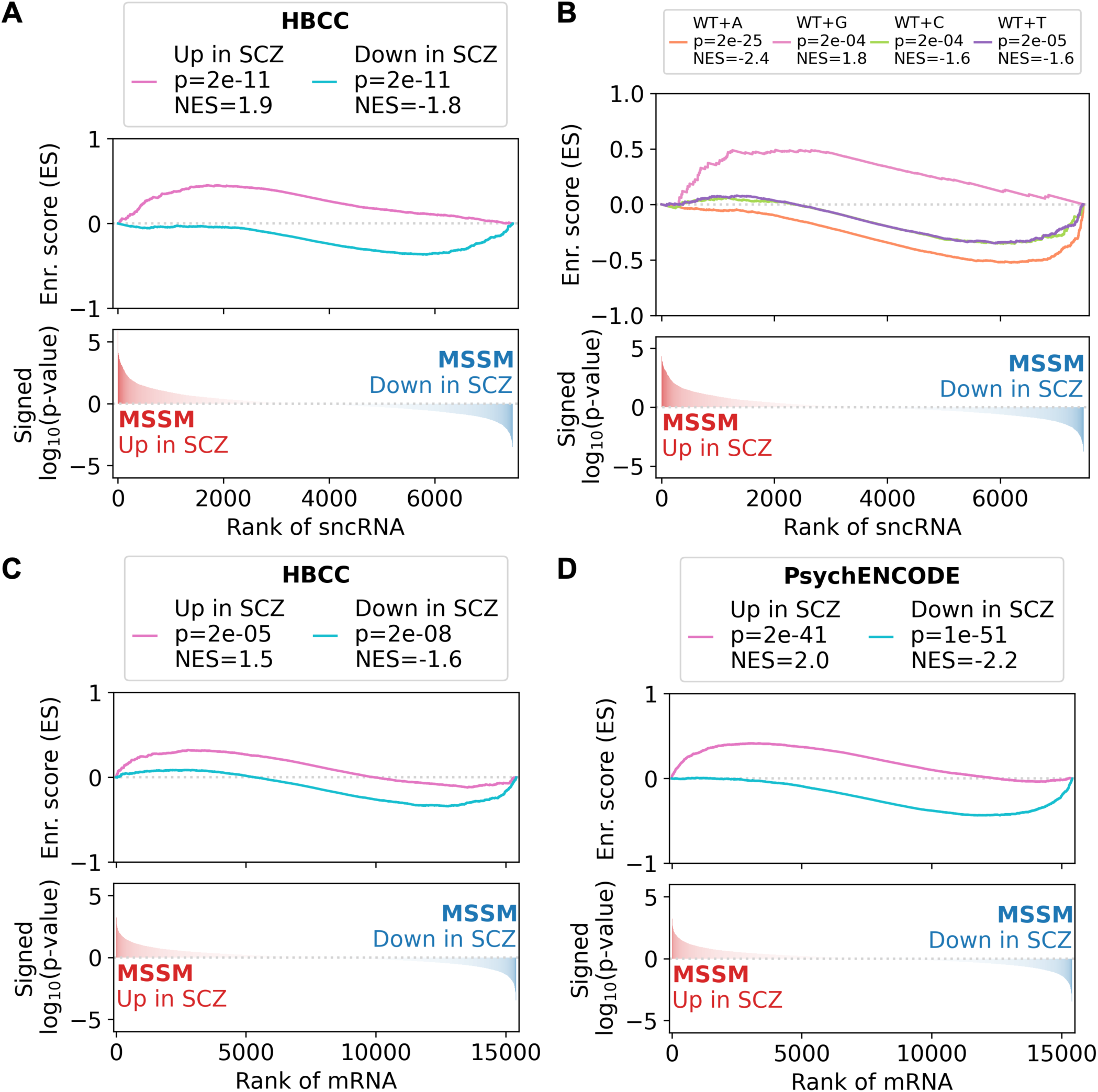
Replication of the HBCC-derived findings in the MSSM brain bank. All abundant sncRNAs (A-B) or mRNAs (C-D) are ranked according to the log_10_ p-value multiplied by the fold change sign of the SCZ vs. control comparison in the MSSM. The following sets were used as references: (A, C) sncRNAs/mRNAs significantly differentially abundant in SCZ (FDR < 0.05) in the HBCC; (B) different non-templated isomiRs; (D) the PsychENCODE set of differentially abundant mRNAs in SCZ.

**Supp. Figure S8.**
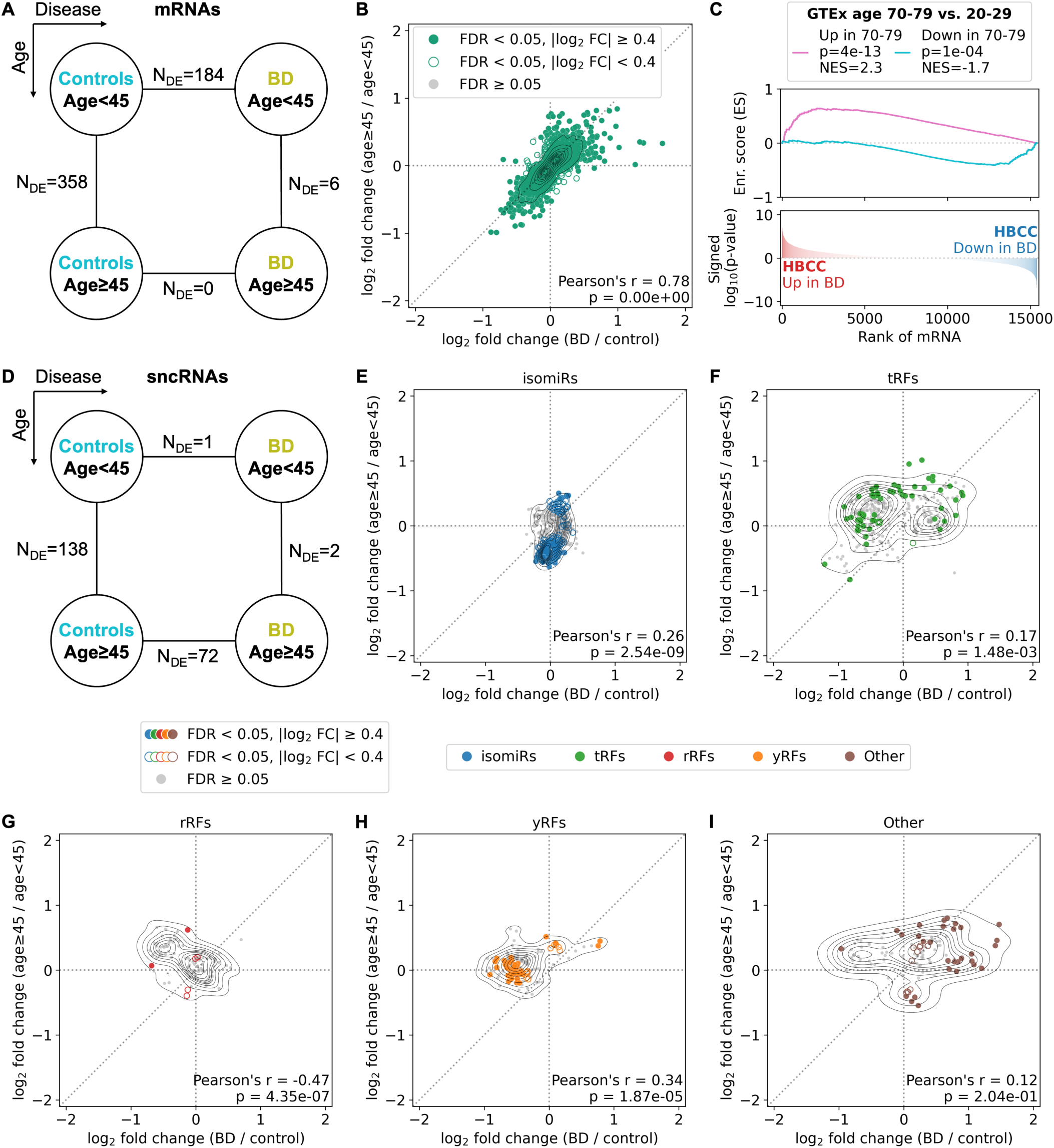
Age-dependent differential mRNA and sncRNA abundance between BD cases and controls (HBCC brain bank). (A, D) The number of significantly differentially abundant mRNAs/sncRNAs (FDR < 0.05) between age-restricted BD vs. control comparisons (horizontal dimension) and condition-restricted age ≥ 45 vs. age < 45 comparisons (vertical dimension). To remove the effect of sample size on the reported numbers, we show median numbers of differentially abundant molecules derived from 100 random down-samplings (see Methods). (B, E-I) Mutual distribution of FCs from BD vs. control and age ≥ 45 vs. age < 45 comparisons. The diagonal dotted line corresponds to equal FCs (Y = X). Only molecules associated with disease status or age group (likelihood-ratio test FDR < 0.05) are shown. Significantly differentially abundant mRNAs/sncRNAs are highlighted with color if FDR < 0.05 in either of the two comparisons. Highlighted markers are filled if |log2 FC| ≥ 0.4 in either of the two comparisons. IsomiRs, tRFs, rRFs, and yRFs include both wild-type and LD ≤ 2 molecules. “Other” molecules include rpFs and additional genomic mappings (see Methods). (C) GSEA of differentially abundant mRNAs in BD in the HBCC brain bank with the GTEx aging signature as a reference gene set.

**Supp. Figure S9.**
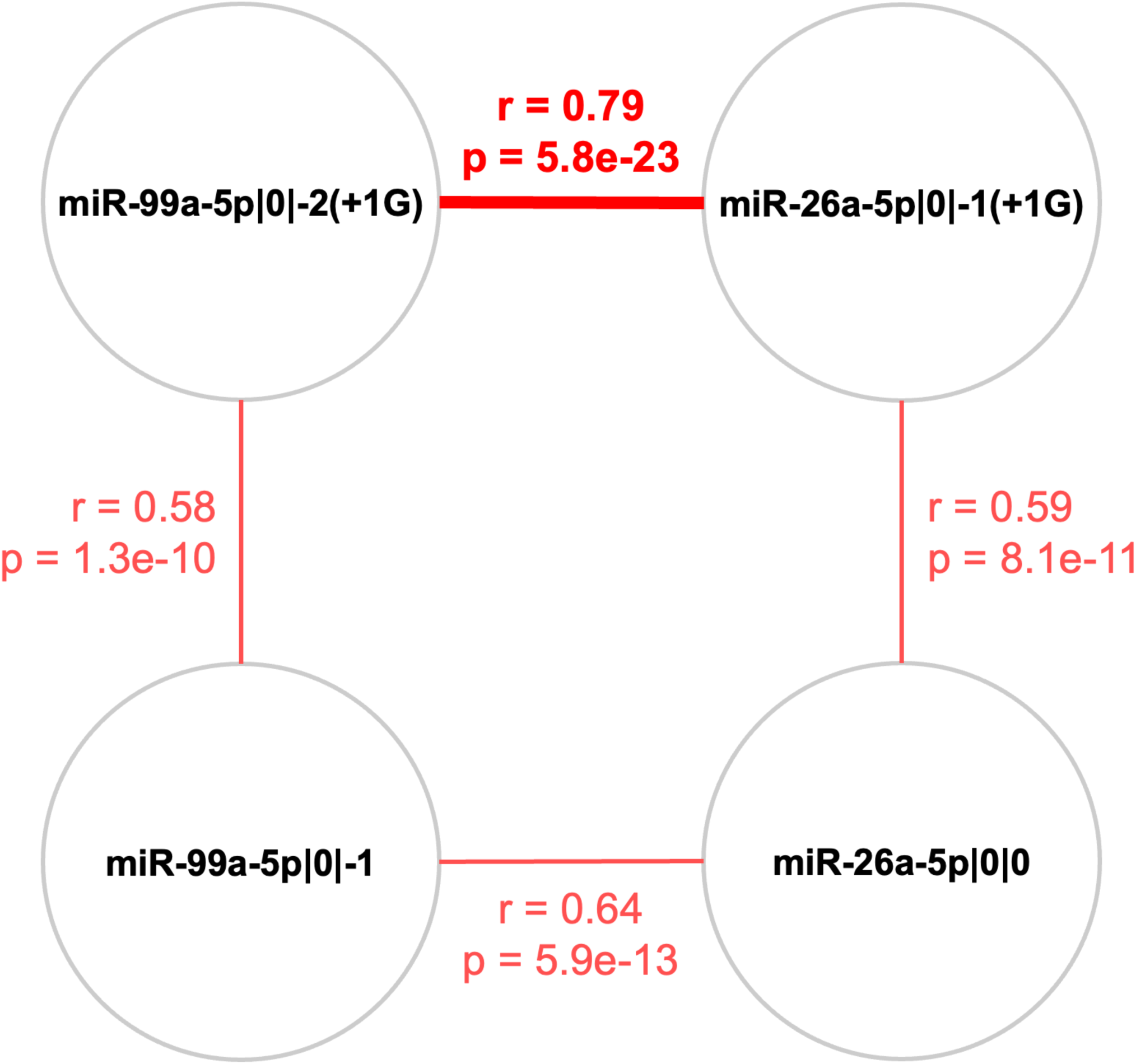
Pearson correlations between different isomiRs of miR-99a-5p and miR-26a-5p. Guanylated isomiRs originating from different miRNAs located on different chromosomes (top row) show higher correlation with each other than with their most abundant sibling isomiRs from the same miRNA (bottom row).

